# Metabolic stress reveals widespread accumulation of cap-unmethylated RNAs

**DOI:** 10.64898/2026.02.23.707474

**Authors:** Zheng Xing, Benjamin M. Sutter, Anna Violet Freitas, Ngoc Khoi Dang, Nicholas T. Ingolia, Benjamin P. Tu

## Abstract

RNA polymerase II transcripts are capped with N^7^-methylguanosine (m⁷G), a conserved modification important for mRNA function. Although traditionally viewed as constitutive, we developed a mass spectrometry-based method to demonstrate that in both yeast and mammalian cells, a substantial population of mRNAs lack cap methylation in response to SAM-limiting conditions and other stress conditions, which may be frequently encountered across organisms. Through developing two transcriptome-wide approaches, we found that methylation is enriched on specific transcripts and uncovered an unexpected connection between histone H3K36me3 and cap methylation, with both marks preferentially associated with stress-responsive MAPK signaling pathways. Strikingly, cap-unmethylated mRNAs exhibit features of canonical mRNAs—they are polyadenylated, exported to the cytosol, and translated. Enforced cap methylation reduces cell growth under SAM limitation, suggesting that unmethylated mRNAs confer an adaptive advantage during stress. These findings establish mRNA cap methylation as a modification responsive to metabolic stress and a previously underappreciated layer of gene expression control.

## Introduction

The metabolism of mRNAs represents a key aspect in the regulation of gene expression and cellular functions. A eukaryotic mRNA consists of several components: a 5’ cap, 5’ and 3’ untranslated regions (UTRs), a coding region, and a 3’ poly(A) tail. Decades of research have established that many of these components are subject to regulatory control. For example, the UTRs and poly(A) tail can vary in length in response to stress conditions^1,2^. Poly(A) tails can undergo deadenylation and re-adenylation as part of the post-transcriptional regulation of maternal mRNAs in the oocyte-to-embryo transition^3^. Introns can be alternatively spliced as a means of gene expression regulation^4^. However, fewer mechanisms have been attributed to regulation of the mRNA cap itself.

Recent years have seen discoveries of non-canonical cap structures derived from metabolites, such as nicotinamide adenine dinucleotide (NAD^+^) and flavin adenine dinucleotide (FAD)^5^. Metabolite caps are not prevalent compared to canonical cap structures (∼1000-fold lower in amount) in human cells^6^. The abundance of such metabolite caps tends to correlate with the cellular levels of the corresponding metabolites^5^, suggesting that mRNA cap composition could change according to the metabolic status of the cell, and possibly alter mRNA metabolism. Intriguingly, the most abundant cap structure in the cell, the m^7^GpppN cap, requires one key metabolite, S-adenosylmethionine (SAM), for its synthesis. The methylation of the mRNA cap has long been recognized as critical for mRNA processing and function, particularly for canonical translation, owing to its specific recognition by the initiation factor eIF4E^7^. SAM is the methyl currency that links environmental conditions to the growth or survival state of the cell^8^. Genes encoding SAM synthetases have been identified as high copy suppressors for a conditional cap methyltransferase (*ABD1*) mutant in yeast^9^, suggesting that cap methylation by Abd1 may be sensitive to SAM levels.

Since its discovery in the 1970s^10^, the m⁷G cap has been considered a constitutive modification. In the last two decades, however, several lines of evidence suggest the existence of cap-unmethylated mRNAs. Overexpression of gene-specific transcription factors c-Myc or E2F1 increases m^7^G-capped RNA levels on several of their target genes^11–13^. Phosphorylation of the mammalian cap methyltransferase (RNMT) was shown to affect the m^7^G-capped levels of three mRNAs tested^14^. However, these studies relied on anti-m⁷G antibodies, which may cross-react with internal m⁷G modifications present in mammalian RNAs^15^. More recently, a mass spectrometry-based method (CAP-MAP) detected unmethylated caps in cells lacking RNMT or RNMT-activating miniprotein (RAM)^16,17^. Despite these studies which suggest RNMT activity could be regulated, transcripts lacking cap methylation are widely believed to be rare and degraded. In line with this, the DXO family of decapping enzymes and exonucleases is proposed to act as a quality control mechanism to clear mRNAs with GpppN unmethylated caps from cells^18–20^. Here, we find that cap-unmethylated transcripts become prevalent under metabolic stress, such as conditions that restrict intracellular amounts of the biological methyl donor SAM. Our results demonstrate that cap methylation is directed to subsets of mRNAs governing stress responses, which can remain functional and play critical roles in cellular adaptation under such conditions.

## Results

### GpppN-capped mRNAs can accumulate under SAM-limiting conditions

We employed a previously established method to induce methionine/SAM restriction in yeast by shifting a prototrophic strain from rich lactate medium (YPL) to synthetic minimal lactate medium (SL)^21^. Such growth conditions enforce respiratory metabolism, which necessitates an increased dependency on sulfur-containing metabolites. To quantitate cap methylation of mRNAs, we digested cellular RNAs and quantified cap dinucleotide structures using liquid chromatography–tandem mass spectrometry (LC-MS/MS) (Fig. 1A, B). Using external standards (Fig. S1A), we quantified cap structures with the first transcribed nucleotide (+1) being a purine, as purines—particularly adenine—are more common than pyrimidines at the +1 position^22^. Unexpectedly, GpppN-capped mRNAs accumulated to ∼50% after 1 hour of SAM restriction (Fig. 1C), while total mRNA levels remained unchanged (Fig. 1D, Fig. S1B). As m^7^GpppA and m^7^GpppG caps behave similarly in our assays, we present only m⁷GpppA cap data in subsequent experiments. To determine whether GpppN-capped mRNAs are polyadenylated, we performed oligo(dT) selection and observed no change in the percentage of GpppA-containing transcripts, indicating that these unmethylated mRNAs possess poly(A) tails (Fig. 1E). This accumulation was further validated by dot blot using an m⁷G-specific antibody with poly(A)+ transcripts (Fig. 1F).

**Fig. 1.**
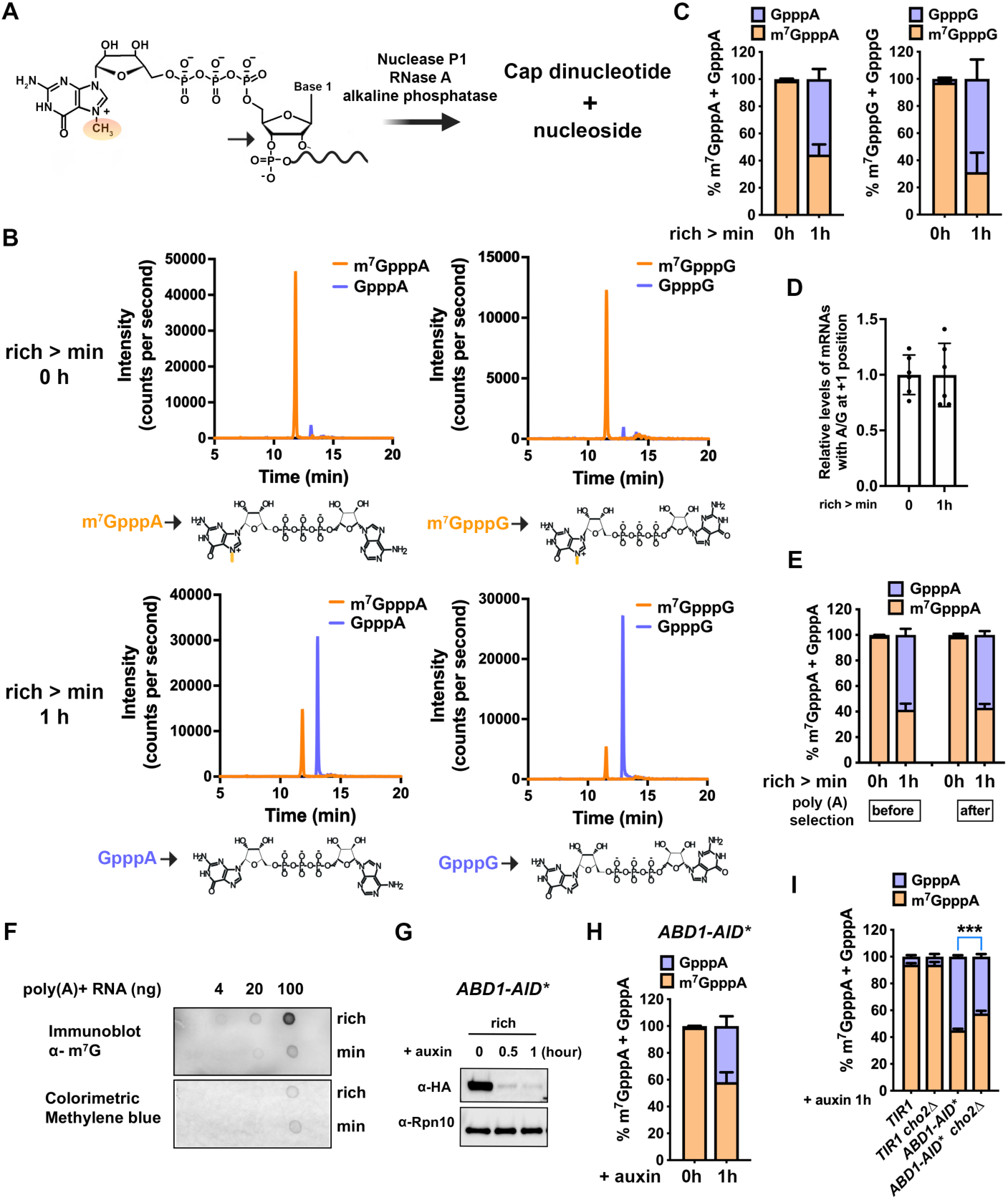
Accumulation of GpppN cap-unmethylated mRNAs in response to methionine and SAM limitation. **(A)** Schematic of mRNA digestion releasing cap dinucleotides and nucleosides. **(B)** LC-MS/MS chromatograms of cap dinucleotides from mRNAs of cells grown in rich (YPL, top) or minimal (SL, bottom) medium for 1 h. Structures of the species representing major peaks are shown below each panel. **(C)** Quantification of cap dinucleotide structures in rich vs. minimal medium using external standards. **(D)** Relative abundance of total mRNAs with A or G at +1 position, measured by LC-MS/MS and normalized to total adenosine. **(E)** GpppA and m⁷GpppA cap percentages before/after oligo(dT) selection. **(F)** Dot blot of poly(A)-selected mRNAs from cells grown in rich or minimal medium, probed with an m⁷G antibody. Methylene blue staining, which binds nucleic acids, was used to assess total RNA loading. **(G)** Western blot showing Abd1-AID*-HA depletion over time with auxin (200 µM). **(H)** Percentage of GpppA or m^7^GpppA cap structures before and after Abd1 depletion for 1 h. **(I)** Percentage of GpppA cap structures of *TIR1*, *TIR1 cho2Δ*, *ABD1-AID\** and *ABD1-AID* cho2Δ* cells, after addition of 200 µM auxin in rich medium for 1 h. All bar graph data represent ≥ 3 biological replicates and are shown as mean ± standard deviation (SD). *** indicates p < 0.001.

The essential protein Abd1 is the sole cap methyltransferase in yeast^23^. Depletion of Abd1 using an auxin-inducible degron (AID*) tag (Fig. 1G)^24^ led to accumulation of GpppA-capped mRNAs (Fig. 1H). This accumulation was partially rescued by increasing cellular SAM levels via deletion of the phosphatidylethanolamine methyltransferase (PEMT) *CHO2*, which was shown to be a major consumer of SAM for phospholipid methylation (Fig. 1I)^25^. Likewise, supplementation of the medium solely with SAM or methionine reversed GpppA mRNA accumulation, whereas supplementation with a non-sulfur amino acid mixture or glucose had no effect (Fig. S1C). Previous work from our laboratory showed that intracellular SAM levels decline most prominently 1 h after the switch to minimal medium and gradually recover by 4 h^26^. Notably, GpppA levels closely mirrored this pattern (Fig. S1D). These findings demonstrate that mRNA cap methylation is sensitive to cellular SAM levels and cap methyltransferase abundance. Importantly, methionine/SAM limitation did not alter Abd1 levels (Fig. S1E), confirming that GpppA mRNA accumulation under SAM limitation is not due to changes in Abd1 expression.

In principle, GpppN-capped mRNAs could arise during new transcription or as a result of demethylation of an m⁷G-capped mRNA. To this end, we tested whether the GpppN mRNAs we observed were newly synthesized. Upon switching cells from rich to minimal medium, we added [^13^C]-labeled guanine (M+3) and readily detected heavy-labeled cap dinucleotides using LC-MS/MS (Fig. S1F), revealing that both m^7^GpppA and GpppA mRNAs can be newly synthesized. Incomplete labeling likely reflects limited incorporation of heavy guanine into guanosine (Fig. S1G). We next added the transcription inhibitor thiolutin when switching cells to minimal medium, which reduced the accumulation of GpppA mRNAs (Fig. S1H), suggesting that they arise from new transcription. This treatment did not completely eliminate GpppA-capped mRNAs, likely because thiolutin may not instantaneously stop all RNA polymerase II transcription^27^. In addition, deletion of individual genes encoding known demethylase enzymes did not alter GpppA accumulation (Fig. S1I), and no eukaryotic enzyme has been identified that specifically removes the N^7^ methyl group from the mRNA cap. Together, these findings support the conclusion that GpppN-capped mRNAs can be generated during new transcription, although the possibility of cap demethylation cannot be excluded.

Rai1 and Dxo1 are yeast decapping and exonuclease enzymes that act on nonconventional mRNA cap structures *in vitro*, including the GpppN cap^19^. They are proposed to serve as quality control factors that clear unmethylated, GpppN-capped mRNAs *in vivo*. To test this, we generated single and double deletion mutants of *RAI1* and *DXO1*. Surprisingly, these deletions did not lead to increased GpppA-capped mRNA accumulation (Fig. S1J). Furthermore, overexpression of both did not change the accumulation of GpppA-capped mRNAs (Fig. S1K, L). Consistent with these findings, Rai1 and Dxo1 protein levels were not induced following the shift to minimal medium (Fig. S1M). Thus, clearance of transcripts with unmethylated caps may not represent the primary role of these enzymes during SAM limitation.

### Cap methylation is enriched for a subset of mRNAs

Having established that mRNA cap methylation is not constitutive, we next investigated whether mRNAs might be differentially methylated. To measure the extent of cap methylation on individual mRNAs, we developed an m^7^G immunoprecipitation (m^7^G-IP) method, utilizing an antibody (K121) that recognizes N^7^-methylated guanosine^18^ to enrich for m⁷G-capped mRNAs from the poly(A)+ RNA pool (Fig. 2A). To validate antibody specificity, we demonstrated that K121 binding to mRNA is competed by methylated, but not unmethylated, cap analogs (Fig. S2A). It also exhibits increased mRNA affinity as compared to IgG control (Fig. S2B), and binds selectively to m⁷G-capped *in vitro* transcribed RNA (Fig. S2C), consistent with previous studies ^18^. We quantified cap methylation by comparing RNA-sequencing read counts before (input) and after (IP) enrichment. By arbitrarily setting cap methylation in rich medium to 100%, we were able to estimate the % methylation for each transcript following 1 hour of methionine/SAM starvation. This revealed that m⁷G is not uniformly distributed across mRNAs, but is instead enriched on a specific subset of transcripts. Meanwhile, a large proportion of transcripts exhibit ∼40–50% cap methylation, consistent with the overall bulk cap methylation level (Fig. 2B, Data S1).

**Fig. 2.**
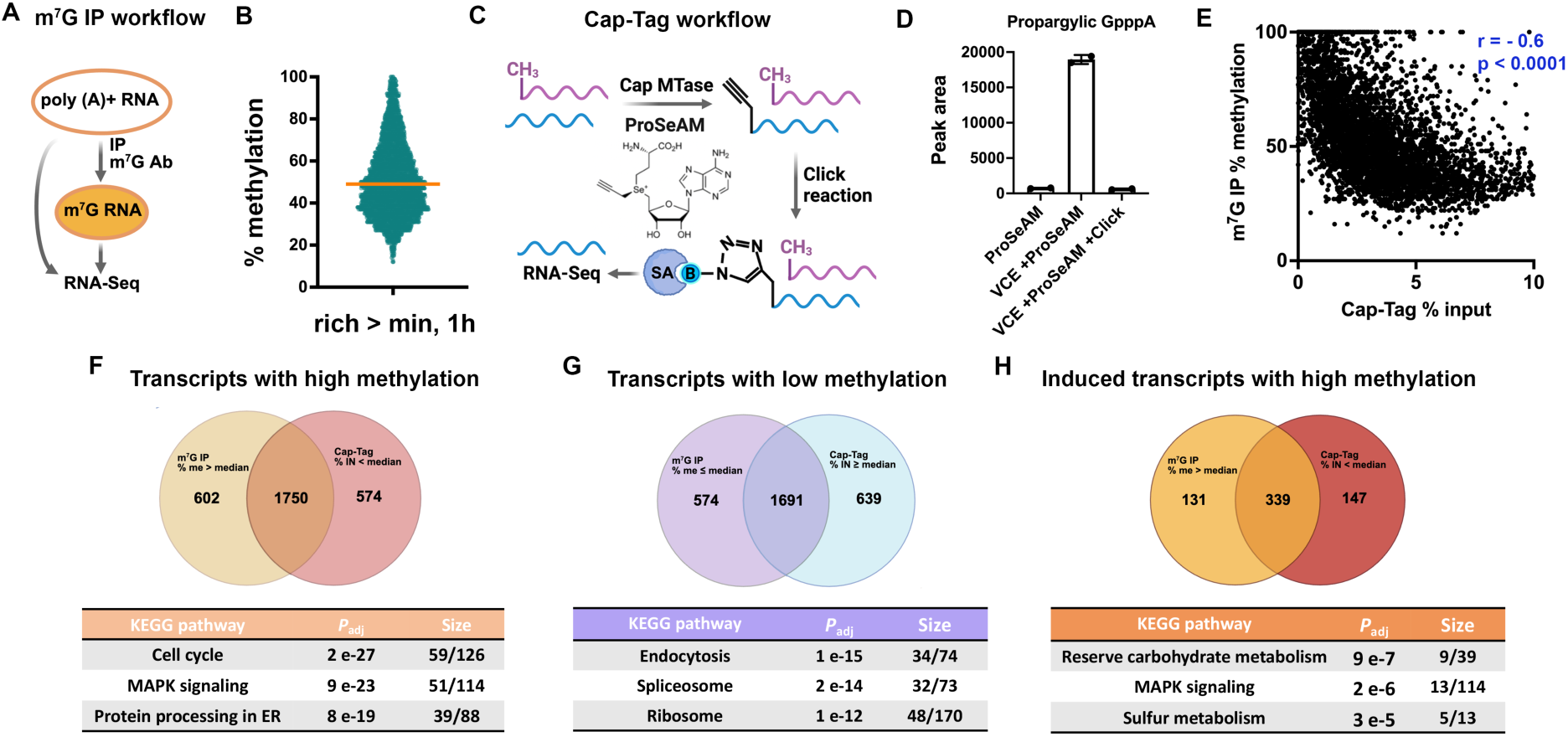
m⁷G cap methylation exhibits transcript-specific distribution. **(A)** Schematic of m^7^G IP workflow. **(B)** Scatter plot showing distribution of mRNA methylation levels after 1 h in minimal medium (methionine/SAM starvation); yellow line indicates median. Transcripts with 100% methylation were omitted from the plot to improve visualization of the distribution but were included in the median calculation. **(C)** Schematic of Cap-Tag workflow. **(D)** LC-MS/MS analysis of propargylic-GpppA after treatment of poly(A)+ RNA with: ProSeAM only (bar 1); Vaccinia capping enzyme (VCE) + ProSeAM (bar 2); and VCE + ProSeAM followed by Click biotinylation, which diminishes propargylic-GpppA (bar 3); n=2. **(E)** Correlation between % methylation (m⁷G-IP) and % input (Cap-Tag); Pearson r was calculated from 4,618 mRNAs. **(F-H)** Venn diagrams showing overlap of high-/low-methylation transcripts identified by both methods, or highly methylated transcripts that are also induced by 2-fold. High-methylation transcripts were defined as those with % methylation > median in m⁷G-IP and % input < median in Cap-Tag; low-methylation transcripts were defined using the opposite criteria. High-confidence (overlapping) transcripts were used for KEGG enrichment analysis via YeastEnrichr^53^. Size shows the numbers of genes in the high-confidence list/number of genes in the KEGG term.

To complement our m⁷G-IP results, we developed an orthogonal method, termed the *in vitro* Cap-Tag assay. Briefly, poly(A)+ RNAs were incubated with a cap methyltransferase and a SAM analog called ProSeAM^28^, which enables incorporation of a propargyl group into GpppN-capped mRNAs, while not affecting already-methylated m^7^GpppN-mRNAs. The propargyl group enables click chemistry-based biotinylation and subsequent enrichment of transcripts that lack cap methylation. Biotinylated mRNAs are captured with streptavidin beads and sequenced (Fig. 2C). LC-MS/MS confirmed propargyl incorporation into GpppA-capped mRNAs, and the click reaction reduced signal intensity, consistent with successful biotin addition (Fig. 2D). *In vitro* transcribed, m^7^GpppG-capped RNA and mRNAs from cells grown in rich medium were tested as negative controls and showed minimal enrichment due to low biotinylation (Fig. S2D, E). Cap-Tag was performed using poly(A)+ RNAs from cells subjected to 1 h of methionine/SAM starvation. Control samples omitting the cap methyltransferase were used to estimate background and subtracted from the final data. We calculated % input to represent GpppN enrichment and compared this to % methylation from m⁷G-IP for each transcript. The two datasets showed strong anticorrelation (Fig. 2E), consistent with the enrichment of methylated mRNAs by m⁷G-IP along with unmethylated mRNAs by Cap-Tag.

We defined high- and low-methylation transcripts using the median value from each dataset and identified overlapping genes as high-confidence targets. A substantial overlap was observed between the two datasets (Fig. 2F-H). Pathway enrichment analysis revealed that dynamic processes such as cell cycle and MAPK signaling are enriched in genes with highly cap-methylated mRNAs (Fig. 2F), and terms such as endocytosis, spliceosome, and ribosome are enriched in genes with lowly methylated mRNAs (Fig. 2G). Although cap methylation levels did not correlate with mRNA abundance or induction (Fig. S2F, G), we sought to exclude the influence of long-lived transcripts synthesized and methylated prior to starvation. To this end, we repeated the pathway enrichment analysis using only genes with induced expression. This analysis revealed enrichment for reserve carbohydrate metabolism, MAPK signaling, and sulfur metabolism (Fig. 2H) - pathways closely associated with nutrient stress responses. Taken together, we have confirmed that cap methylation is selectively enriched in a subset of mRNAs during methionine/SAM starvation.

### Cap methylation exhibits correlation with elevated H3K36me3 levels, and is linked to MAPK signaling

How might differential methylation of individual mRNAs be achieved? Abd1 is thought to methylate the mRNA cap co-transcriptionally by binding to the C-terminal domain of RNA polymerase II (Pol II)^29^. We performed chromatin immunoprecipitation followed by sequencing (ChIP-seq) against Abd1-HA and Rpb1, a subunit of the RNA Pol II holoenzyme, using cells grown in minimal medium for 1 h. Consistent with prior findings^29^, Abd1 signal peaks at the 5′ ends of genes (Fig. S2H) and shows a strong genome-wide correlation with Pol II occupancy (Fig. S2I). To assess Abd1 binding without influence by transcriptional activities, we quantified relative Abd1 occupancy as the ratio of Abd1/Pol II signal. This metric shows a weak to moderate correlation with the percentage of cap methylation (Fig. S2J). We then repeated this analysis in cells grown in rich medium. Approximately half of the Abd1 peaks identified under SAM limitation overlap with those detected in rich medium (Fig. S2K, Data S2), consistent with extensive transcriptional reprogramming between the two conditions. However, Abd1 occupancy at these shared peaks is highly correlated (Fig. S2L), suggesting that differential Abd1 recruitment is unlikely to account for transcript-specific differences in mRNA cap methylation.

We previously demonstrated that bulk histone methylation is responsive to cellular SAM levels, with global histone methylation—and particularly H3K36me3—reduced upon SAM limitation^26^. To investigate whether mRNA cap methylation might correlate with histone methylation, we performed ChIP-seq for H3K4me3, H3K36me3, and H3K79me3. As expected, these marks show canonical distributions—H3K4me3 enriched at transcription start sites (TSSs) and H3K36me3/H3K79me3 across gene bodies. Consistent with earlier results, we observed the most pronounced global reduction in H3K36me3 upon starvation (Fig. 3A, Fig. S3A, B, and Data S2). Surprisingly, a subset of genes displayed a marked increase in H3K36me3 following methionine/SAM limitation (Fig. 3B). Changes in H3K36me3 levels showed very little correlation with changes in mRNA expression levels (r = 0.08, Fig. S3C), consistent with the notion that H3K36me3 may also serve a metabolic function that can supersede its role in transcriptional regulation^25,26,30^. Interestingly, elevated H3K36me3 strongly predicts high levels of cap methylation (Fig. 3C). Consistent with this observation, strains lacking the H3K36 methyltransferase *SET2*^31^ or harboring the non-methylatable *H3K36A* mutation displayed reduced m⁷GpppA levels (Fig. 3D), suggesting that H3K36me3 enhances cap methylation for a subset of transcripts.

**Fig. 3.**
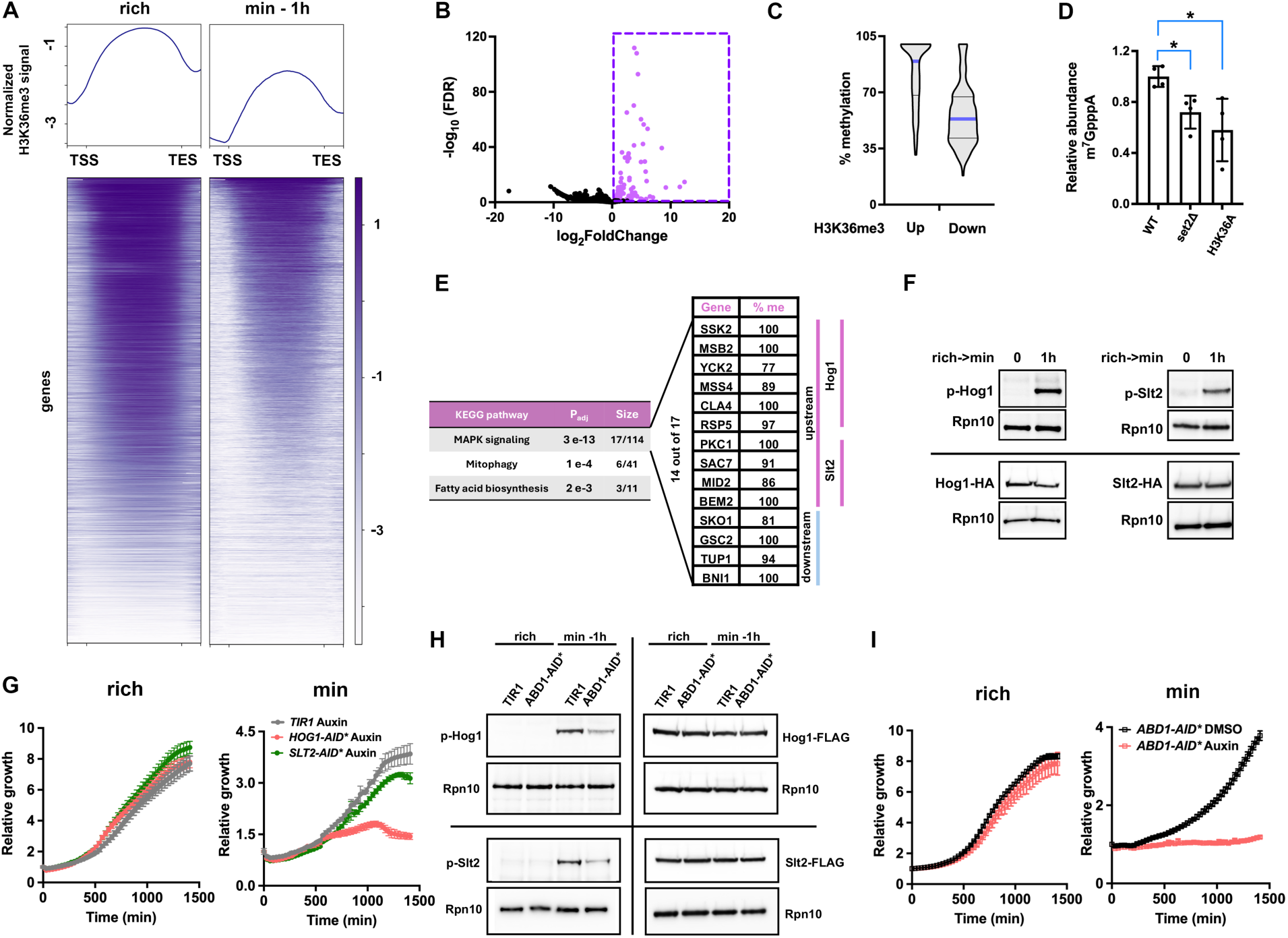
Selective mRNA cap methylation associates with H3K36me3 levels. **(A)** H3K36me3 ChIP-seq profiles in rich or minimal medium, normalized to total H3. **(B)** Volcano plot highlighting 227 genes (magenta) with increased H3K36me3 after 1 h in minimal medium (log₂FC > 0, FDR < 0.05). **(C)** Violin plots comparing percent cap methylation in genes with upregulated H3K36me3 versus genes with downregulated H3K36me3. **(D)** Relative abundance of m^7^GpppA cap structures in wild type, *set2Δ,* and *H3K36A* mutant after 1 h in minimal medium; mean ± SD, n = 4. * indicates *p* < 0.05. **(E)** KEGG pathways enriched among genes with increased H3K36me3 upon methionine starvation. Size shows (H3K36me3-upregulated genes)/(total genes in pathway). Notably, 14 out of the 17 genes in the MAPK pathway exhibit high cap methylation. **(F)** Western blot of Hog1 and Slt2 phosphorylation after 1 h in minimal medium; separate HA blots show total Hog1 and Slt2 protein levels remain unchanged. **(G)** Growth curves of *TIR1* and Hog1 or Slt2-depleted cells in rich or minimal media containing 40 µM of auxin; mean ± SD, n = 6. **(H)** Western blot showing Hog1 and Slt2 phosphorylation after 1 h in minimal medium in both *TIR1* and Abd1-depleted strains; FLAG-blots show total Hog1 and Slt2 protein levels remain unchanged after the media switch. **(I)** Growth curves of the *ABD1-AID\** strain treated with DMSO or 40 µM auxin to achieve partial Abd1 depletion in rich or minimal medium. Data are presented as mean ± SD (*n* = 3).

Pathway analysis of these genes with upregulated H3K36me3 identified MAPK signaling as the top enriched pathway. Among 17 MAPK pathway genes with elevated H3K36me3, 14 also exhibited high cap methylation (Fig. 3E). Examination of these highly cap-methylated transcripts revealed enrichment for regulators in the Hog1 and Slt2 MAPK branches, with 10 genes positioned upstream of Hog1 or Slt2. In line with this, Hog1 and Slt2 were phosphorylated upon switching to minimal medium (Fig. 3F). Moreover, depletion of these kinases—especially Hog1— compromised growth in minimal but not rich medium (Fig. S3D and Fig. 3G). Finally, disruption of cap methylation by depleting Abd1 reduced Hog1 and Slt2 phosphorylation after the media switch without affecting their total protein levels (Fig. 3H), supporting a role for prioritized cap methylation in promoting MAPK activation. In line with this, partial degradation of the essential protein Abd1 resulted in more pronounced growth defects in minimal medium than in rich medium (Fig. 3I). Together, these findings indicate that both H3K36me3 and mRNA cap methylation respond similarly to SAM starvation: they are globally downregulated but selectively maintained or elevated at a shared subset of genes and their encoded transcripts, such as those involved in MAPK signaling—a pathway critical for stress adaptation.

We next examined cap methylation of transcripts encoding the upstream MAPK regulators following Abd1 depletion. As expected, cap methylation was reduced for all mRNAs analyzed, with the most pronounced reductions observed for *MID2*, *MSB2*, and *SSK2*, even after accounting for changes in mRNA abundance (Fig. S3E, F). Western blot analysis further showed that Ssk2, a MAPK kinase kinase (MAPKKK)^32^, exhibited the greatest reduction in protein levels (Fig. S3G). To determine which of these upstream regulators is most critical for MAPK activation, we individually deleted seven non-essential genes and examined Hog1 and Slt2 phosphorylation. Among these mutants, ssk2Δ displayed the most pronounced reduction in MAPK phosphorylation (Fig. S3H, I). Taken together, these results suggest that reduced SSK2 cap methylation and the accompanying decrease in Ssk2 protein abundance may contribute to the reduced MAPK activation observed upon Abd1 depletion. These data provide an example of how cap methylation can influence protein abundance and consequently, cellular phenotypes.

### GpppN mRNAs are exported to the cytosol and can be translated

What is the fate of mRNAs lacking cap methylation? We next sought to characterize GpppN-capped mRNAs, which represent a largely underappreciated class of mRNAs in the cell. The observation that mRNAs are not uniformly methylated and that cap-unmethylated mRNAs are not rapidly degraded suggests they may be functional. To investigate their fate, we first examined the localization of cap-unmethylated mRNAs. We inserted MS2 binding sites (MBSV6^33^) into the 3’ UTR of the *MET16* mRNA, a lowly methylated transcript during SAM starvation (% methylation ∼ 26%) (Fig. 4A). This enabled visualization of *MET16* mRNA via expression of MS2 coat protein (MCP)-GFP. Interestingly, *MET16* mRNA appears as cytoplasmic foci in live cells (Fig. 4B), a finding corroborated by FISH in fixed cells (Fig. S4A). As stress granules and processing-bodies (P-bodies) are common ribonucleoprotein (RNP)-containing foci, we assessed their formation under methionine/SAM limitation. Stress granule markers readily formed foci following heat shock (Fig. S4B) but not under SAM-limiting conditions (Fig. S4C, E). In contrast, P-bodies readily formed under SAM limitation (Fig. S4D). Despite the formation of P-bodies, *MET16* mRNA foci did not colocalize with them (Fig. 4C). Similarly, two additional lowly methylated mRNAs, *BET1* and *RPN12* (approximately 33% and 29% cap methylation, respectively), also formed distinct cytoplasmic foci (Fig. S4F-H). These results suggest that cap-unmethylated mRNAs are exported to the cytoplasm and do not localize to known RNP granules.

**Fig. 4.**
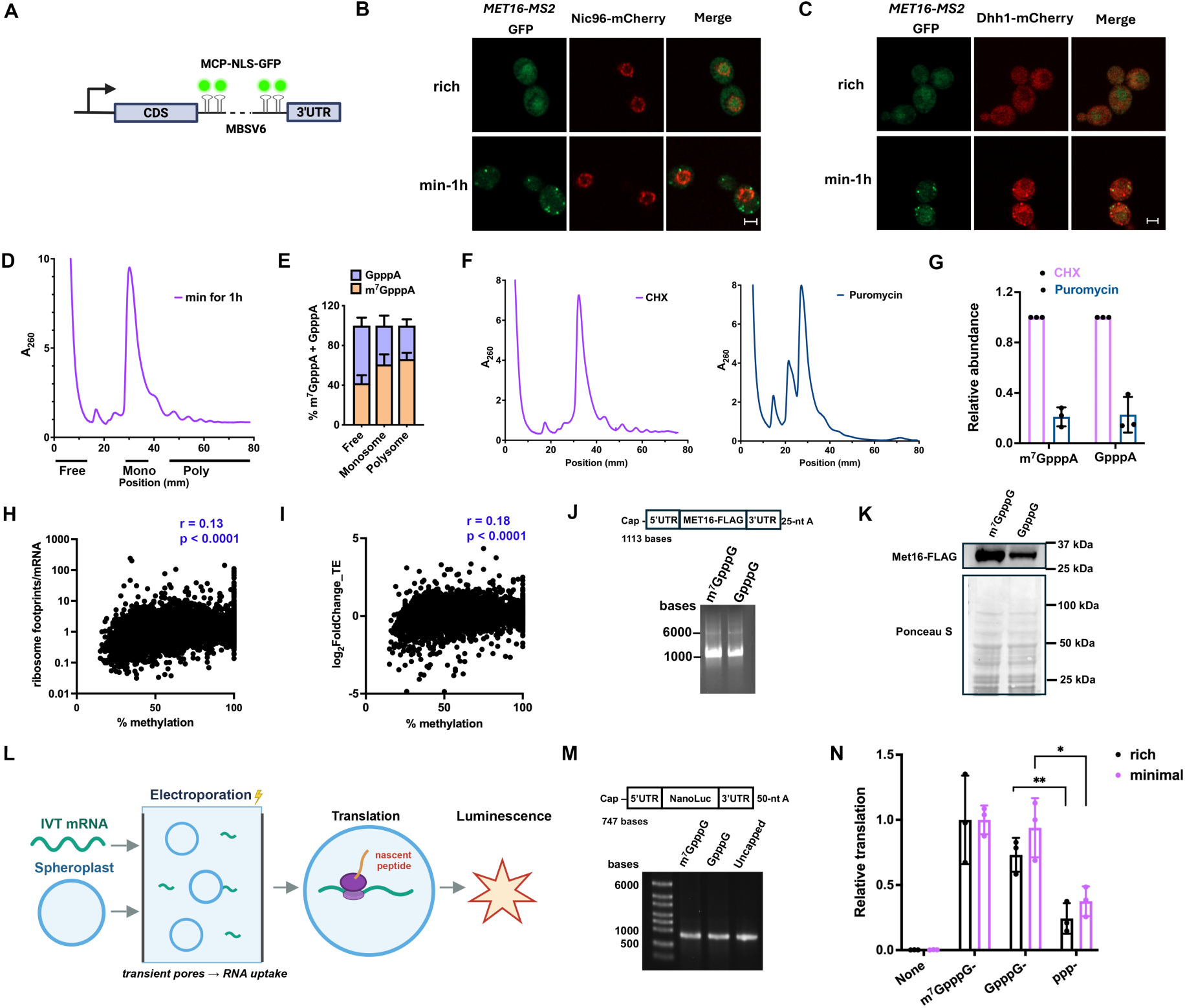
GpppN cap-unmethylated mRNAs can be translated. **(A)** Schematic of MS2-based visualization of an mRNA of interest. **(B)** Live confocal images of MET16 mRNA (green) and nuclear pores (Nic96-mCherry, red); scale bar, 2 µm. **(C)** Live confocal images of MET16 mRNA (green) and P-body marker Dhh1-mCherry (red); scale bar, 2 µm. **(D)** Polysome profiling of cells grown in minimal medium for 1 h. **(E)** LC-MS/MS quantification of GpppA and m⁷GpppA in pooled fractions from (D). **(F)** Polysome profiling of cells in minimal medium (1 h) with CHX (0.1 mg/mL) or puromycin (2 mM) treatment. **(G)** Relative abundance of m⁷GpppA and GpppA after CHX (set to 1) or puromycin treatment, normalized to adenosine. **(H)** Correlation of ribosome footprint/mRNA reads (TE) and methylation levels of each mRNA in cells grown in minimal medium for 1 h. Pearson r = 0.13. **(I)** Correlation of log2FoldChange (FC) of TE (min 1h/rich) and % methylation levels of each mRNA following growth in minimal medium for 1 h. Pearson r = 0.18. **(J)** RNA gel electrophoresis of *in vitro* transcribed *MET16* RNAs with either m^7^GpppG or GpppG cap. **(K)** Western blot analysis of *in vitro* translation of *MET16* RNAs using cell-free yeast translation extracts. Ponceau S stain confirms equal loading of yeast extracts. **(L)** Schematic of the *in vivo* translation assay using electroporation of *in vitro* transcribed mRNAs into yeast. **(M)** RNA gel electrophoresis of *in vitro* transcribed NanoLuc RNAs bearing m^7^GpppG, GpppG, or a triphosphate (ppp, uncapped) at the 5’ end. **(N)** Relative translation of differently capped NanoLuc reporter mRNAs in the *in vivo* translation assay, with m⁷GpppG-capped mRNA recovered in rich medium set to 1. Different RNAs were electroporated into cells in the same buffer, and luciferase activity was normalized to total protein concentration. All bar graph data represent ≥ 3 biological replicates and are shown as mean ± SD. *, ** indicate p < 0.05 and p < 0.01, respectively.

We next asked if cap-unmethylated mRNAs have the potential to be translated. Although global translation is reduced in cells grown in minimal medium compared with rich medium, with an ∼40% decrease in polysome-associated mRNAs (Fig. S5A, B), cells must still synthesize new proteins to adapt to methionine/SAM limitation. We performed polysome profiling on cells grown in minimal medium for 1 h, extracted RNA from fractions representing the free mRNA, monosome, and polysome populations (Fig. 4D), and analyzed cap structures by LC-MS/MS. Surprisingly, we observed that GpppA mRNAs are present in both the monosome and polysome fractions (Fig. 4E). Northern blot analysis confirmed the presence of the lowly methylated *MET16* mRNA in these fractions (Fig. S5C). Furthermore, we treated cell lysates with puromycin to dissociate mRNAs from the polysome fractions (Fig. 4F)^34^. This treatment reduced the polysome association of both m⁷GpppA and GpppA mRNAs to a similar extent (Fig. 4G), validating that GpppA mRNAs associate with active polysomes.

To investigate the relationship between cap methylation and translation efficiency (TE), we performed ribosome profiling on cells collected before and after the switch to minimal medium. TE was calculated as the ratio of ribosome footprint reads to mRNA reads. This analysis revealed a weak positive correlation between TE in minimal medium and mRNA cap methylation levels (correlation coefficient r = 0.13; Fig. 4H), as well as between changes in TE following starvation and cap methylation levels (r = 0.18; Fig. 4I, Data S3). In addition, changes in protein abundance showed essentially no correlation with cap-methylation levels (r = 0.03, Fig. S5D, Data S4). Together, these results support the conclusion that cap-unmethylated mRNAs can be translated, while the weak positive correlation between TE and cap methylation indicates that cap methylation confers a small advantage in translational efficiency globally. To directly test the effect of cap methylation on translation, we *in vitro* transcribed polyadenylated *MET16* RNA containing its native UTRs and coding region, along with a FLAG tag sequence. Transcripts were capped with either m⁷GpppG or GpppG (Fig. 4J) and subjected to *in vitro* translation using yeast cell-free extracts containing the necessary machinery for protein synthesis^35^. Western blot analysis showed that GpppG-capped *MET16* RNA was translated, albeit with lower efficiency than m⁷GpppG-capped RNA (Fig. 4K).

To our knowledge, direct *in vivo* comparisons of translation from otherwise identical mRNAs bearing either an m⁷GpppN or GpppN cap have been reported only in *Xenopus* oocytes during meiotic maturation^36^. In that study, GpppN-capped mRNAs were translated, although less efficiently than their m⁷GpppN-capped counterparts. Whether cap-unmethylated mRNAs can be translated *in vivo* in yeast, however, has not been tested. We therefore sought to directly assess the translational capacity of cap-unmethylated mRNAs in yeast cells. To address this, we generated yeast spheroplasts and introduced *in vitro*-transcribed reporter mRNAs bearing either an m⁷GpppG cap, a GpppG cap, or a 5′-triphosphate via electroporation. The spheroplasts were then recovered in either rich or minimal medium to permit translation (Fig. 4L). To facilitate quantification, we replaced the *MET16* coding sequence with a luciferase reporter while preserving the native *MET16* UTRs (Fig. 4M). GpppG-capped reporter mRNAs produced luciferase signals comparable to those of m⁷GpppG-capped mRNAs in both rich and minimal media, and substantially higher than those of 5′-triphosphorylated (pppG) RNAs (Fig. 4N). Interestingly, we did not observe a significant difference between rich and minimal media. One possible explanation is the use of the *MET16* UTRs, as endogenous *MET16* mRNA retains relatively high translational efficiency (Fig. S6C) despite exhibiting low cap methylation under methionine-limited conditions. Together, these results demonstrate that GpppG-capped mRNAs can directly be translated in living yeast cells.

### GpppN mRNAs are critical during SAM starvation and oxidative stress

Switching yeast cells from rich to minimal medium induces acute methionine/SAM starvation. However, yeast cells can recover from this stress by reprogramming their transcription and metabolism, allowing synthesis of methionine/SAM *de novo* using inorganic sulfur^8^. This transition represents a critical period when the cells undergo active adaptation. To test whether the accumulation of GpppN-capped mRNAs might be beneficial during this phase, we overexpressed *ABD1* under the constitutive *TEF1* promoter. Expectedly, GpppA cap levels are reduced upon Abd1 overexpression (Fig. 5A). However, this impaired the growth of cells in conditions of methionine starvation, but not in methionine/SAM-replete conditions (Fig. 5B), which was not due to a change in cellular methylation potential, as SAM/SAH levels remained unchanged (Fig. 5C). To determine which transcripts are affected by enforced cap methylation, we performed m^7^G-IP on the *pTEF1-ABD1-HA* strain following 1 h of growth in minimal medium. As expected, Abd1 overexpression resulted in a global increase in mRNA cap methylation (Fig. 5D, Data S5). KEGG pathway analysis of transcripts exhibiting more than a 2-fold increase in cap methylation identified ribosome, spliceosome, and endocytosis as the three most enriched pathways (Fig. 5E), mirroring the pathways with low cap methylation in Fig. 2G. These findings suggest that Abd1 overexpression primarily increases cap methylation globally rather than selectively targeting specific transcripts, consistent with our observation that differential Abd1 recruitment is not a major determinant of preferential cap methylation.

**Fig. 5.**
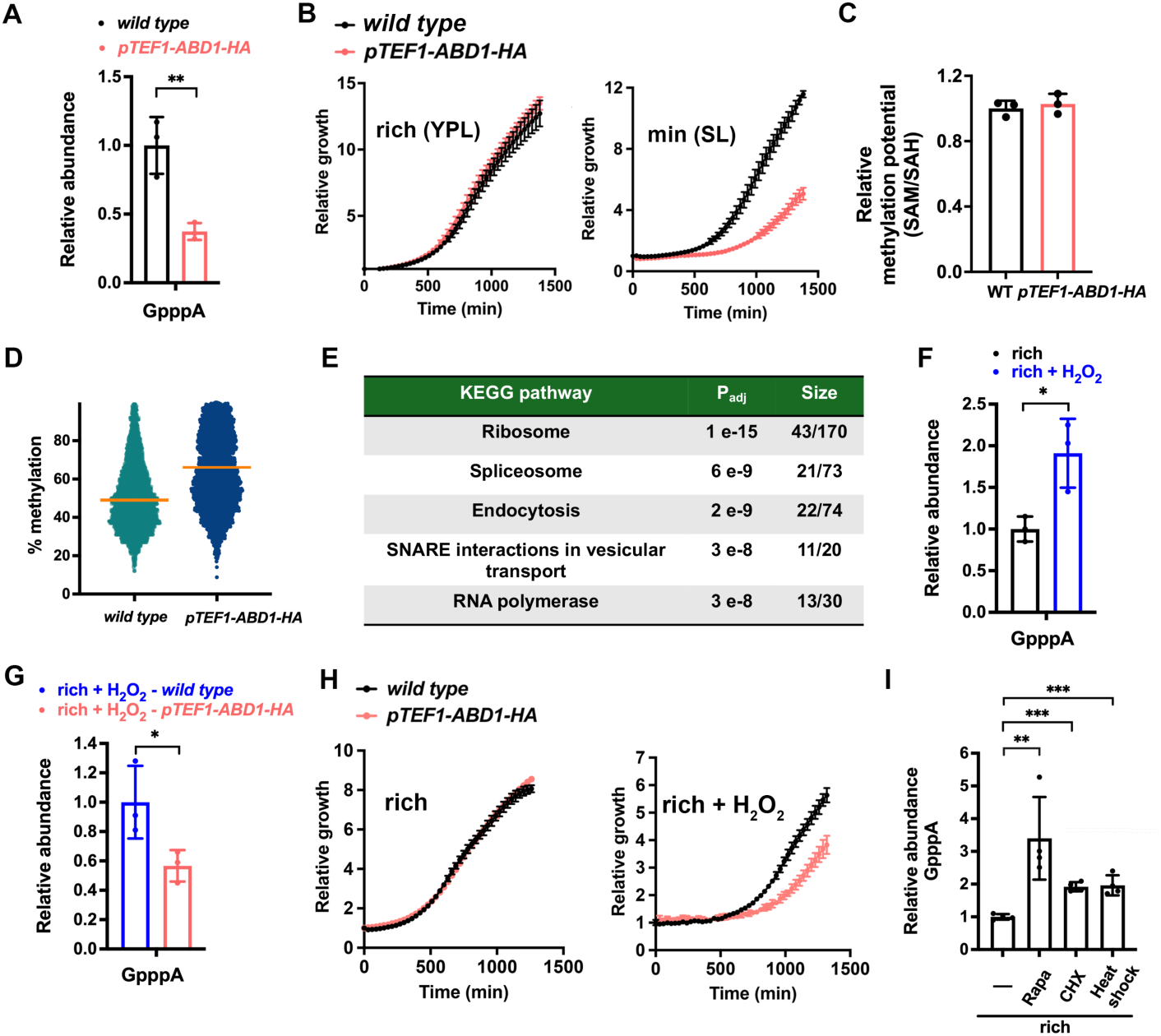
GpppN cap-unmethylated mRNAs are functional. **(A)** LC-MS/MS analysis of relative GpppA levels in minimal medium (1 h) for wild type and *ABD1* overexpression strains. **(B)** Growth curves of wild type and *ABD1* overexpression strains. **(C)** Relative methylation potential (SAM/SAH ratio) in wild-type and *ABD1* overexpression strains. **(D)** Scatter plot showing transcript-specific cap methylation measured by m^7^G-IP in wild-type (Fig. 2B) and *pTEF1-ABD1-HA* strains grown in minimal medium for 1 h. Orange lines indicate the median cap methylation (wild type, 49%; *pTEF1-ABD1-HA*, 66%). Transcripts with 100% cap methylation were omitted from the plot to improve visualization but were included in the median calculation. **(E)** KEGG pathway analysis of genes with a >2-fold increase in cap methylation in *pTEF1-ABD1-HA* relative to wild type. **(F)** LC-MS/MS analysis of relative GpppA levels in rich medium ± 1 mM H₂O₂ (1 h). **(G)** LC-MS/MS analysis of relative GpppA levels of wild type and *ABD1* overexpressing cells grown in rich medium + 1 mM H_2_O_2_ for 1 h. **(H)** Growth curves of wild type and *ABD1* overexpressing cells. **(I)** LC-MS/MS analysis of relative GpppA mRNA levels in cells grown in rich medium, or with addition of 20 ng/mL rapamycin, 25 µg/mL CHX, or 42°C for 1 h. All bar graph data represent ≥ 3 biological replicates and are shown as mean ± SD. *, **, and *** indicate p < 0.05, p < 0.01, and p < 0.001, respectively.

To test if the growth defect is a general consequence of increased cap methyltransferase activity, we expressed the SARS-CoV-2 cap methyltransferase NSP14 in yeast. NSP14 expression reduced growth in minimal medium but not in rich medium (Fig. S5E–G) and decreased GpppA-mRNA levels, consistent with its ability to methylate host mRNA caps in yeast cells (Fig. S5H)^37^. This raises the possibility that viral infection could alter cellular mRNA cap methylation status, representing a previously unrecognized consequence of viral infection.

To explore conditions beyond methionine starvation that might promote the accumulation of cap-unmethylated mRNAs, we treated cells with H₂O₂, an oxidant that may deplete cellular reducing equivalents and SAM^8^, and found that H₂O₂ increased GpppA-mRNA levels (Fig. 5F). Overexpression of Abd1 during H₂O₂ exposure reduced GpppA accumulation (Fig. 5G), which was accompanied by reduced cell growth (Fig. 5H). Additional stress conditions, including TORC1 inhibition by rapamycin, translation inhibition by cycloheximide, and heat shock, also increased GpppA accumulation (Fig. 5I). These findings suggest that the accumulation of cap-unmethylated mRNAs could be a response to multiple forms of cellular stress as a means of stress adaptation.

### Conservation of dynamic mRNA cap methylation in mammalian cells

Mammalian mRNAs also harbor a conserved m⁷G cap, along with additional modifications, including 2′-O-methylation on the first transcribed nucleotide and N⁶-methylation when the first transcribed nucleotide is adenosine^38^. To determine whether the accumulation of unmethylated mRNA caps also occurs in mammalian cells, we subjected human HEK293T cells and mouse Neuro-2a (N2a) cells to methionine starvation^39^. This reduced the abundance of fully methylated cap dinucleotides, while increasing incompletely methylated cap species, including the completely unmethylated GpppN cap (Fig. 6A, B). Because authentic standards are not available for all possible cap dinucleotide species, the relative proportions of individual cap structures within the total cap pool could not be determined.

**Fig. 6.**
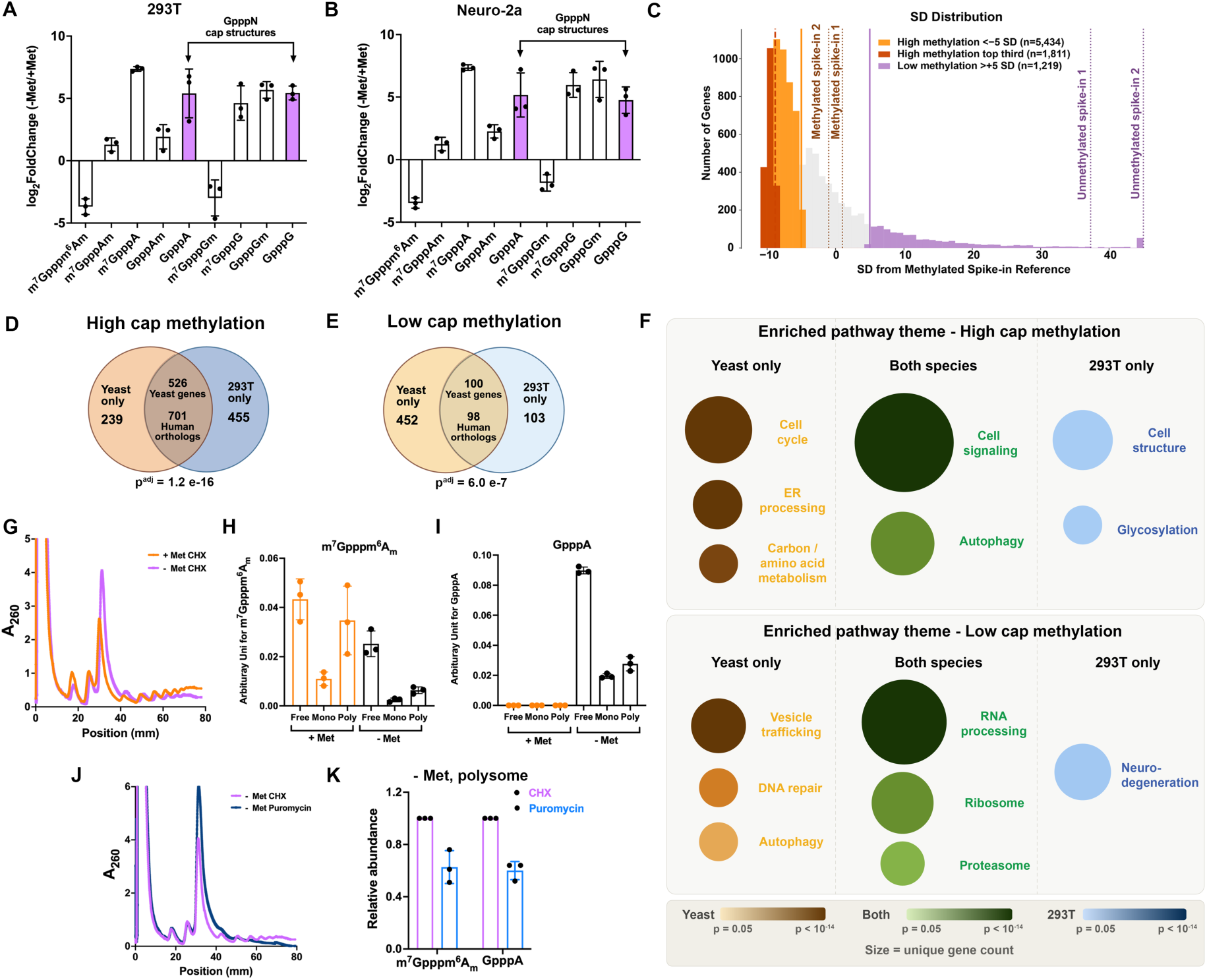
Similar classes of cap-unmethylated transcripts accumulate in mammalian cells in response to methionine starvation. (**A, B)** LC-MS/MS analysis of the relative abundance of cap structures normalized to guanosine in HEK293T (A) and Neuro-2a (B) cells cultured in methionine-replete (+Met) or methionine-depleted (- Met) medium for 18 h. **(C)** Genome-wide mRNA cap methylation analysis using Cap-Tag in 293T cells grown in - Met medium for 18 h. Histogram of mean % input from 10,066 mRNAs expressed as standard deviations (SD) from the methylated spike-in reference. Dashed lines represent the 2 methylated spike-in RNAs (orange) and 2 unmethylated spike-in RNAs (purple). **(D–E)** Venn diagrams showing the overlap of conserved yeast–human orthologs in the high-(D) and low-methylation (E) categories. Analysis was restricted to the 1,817 yeast genes with annotated human orthologs retrieved from Ensembl BioMart (www.ensembl.org) that were also expressed in both the yeast and HEK293T datasets. Orthology relationships included one-to-one, one-to-many, and many-to-many mappings (Data S6). **(F)** Bubble chart summarizing KEGG pathway enrichment themes derived from the 10 most significantly enriched pathways from each of four gene sets: yeast high cap methylation, yeast low cap methylation, 293T high cap methylation (top third), and 293T low cap methylation. Bubble size is proportional to the number of unique genes assigned to each functional theme. Color represents combined p-value calculated using the Cauchy test^54^. **(G)** Polysome profiling of 293T cells cultured in + Met or - Met medium for 18 h and treated with cycloheximide (CHX). **(H–I)** LC–MS/MS quantification of m^7^Gpppm^6^Am (H) and GpppA (I) in RNA isolated from pooled fractions of the polysome profiles. Arbitrary units represent the peak area of each cap dinucleotide normalized to the peak area of adenosine. Free indicates early fractions containing free mRNA and fractions containing some ribosomal subunits. Mono indicates monosome fractions. Poly indicates polysome fractions. **(J)** Polysome profiling of 293T cells cultured in methionine-depleted (- Met) medium for 18 h and treated with cycloheximide (CHX; 0.1 mg/mL, 10 min) or puromycin (0.1 mg/mL, 15 min). **(K)** Relative abundance of m^7^Gpppm^6^Am and GpppA following CHX (set to 1) or puromycin treatment. Cap dinucleotide levels were normalized to adenosine and are presented relative to the CHX-treated condition. All bar graph data represent ≥ 3 biological replicates and are shown as mean ± SD.

We next sought to determine cap methylation amounts for individual mRNAs in mammalian cells. In addition to the 5′ cap, m⁷G can occur internally within mRNAs^15^. While internal m⁷G is absent in *S. cerevisiae*^40^, it has been reported in mammalian cells^15^. Consequently, the m⁷G-IP approach is not suitable for measuring cap methylation in mammalian cells. We therefore adapted our Cap-Tag method, which enriches for cap-unmethylated mRNAs, for use in mammalian cells and applied it to methionine-starved HEK293T cells. To establish the relationship between % input and N^7^-cap methylation status, we included two N^7^-methylated and two unmethylated synthetic spike-in RNAs as internal controls. For each transcript, % input was calculated and the mean and standard deviation (SD) of the values from the two methylated spike-in controls were used as a reference. Transcripts with % input more than 5 SD below the methylated spike-in mean were classified as highly methylated, whereas transcripts more than 5 SD above the methylated spike-in mean were classified as lowly methylated (Fig. 6C, Data S6). To investigate whether transcript-specific patterns of cap methylation are conserved between yeast and mammals, we retrieved all yeast genes with one or more annotated human orthologs from Ensembl. Among these, 1,817 yeast genes and their corresponding human orthologs were expressed in both the yeast and HEK293T datasets. Despite approximately one billion years of evolutionary divergence, we observed a significant overlap between yeast and human orthologs classified as highly or lowly methylated (Fig. 6D, E and Data S6). Conserved genes were enriched for KEGG pathways (Fig. S5I, J) that closely resembled those identified in yeast (Fig. 2F, G).

Because orthology-based analysis was limited to 1,817 genes, we next performed pathway enrichment analyses using the lowly methylated transcripts and the top one-third of highly methylated transcripts identified by Cap-Tag in HEK293T cells and compared the enriched pathways with those identified in yeast (Data S6). This revealed that yeast and mammals share similar functional themes associated with cap methylation status (Fig. 6F). Highly methylated transcripts were enriched in pathways related to cell signaling and autophagy, whereas lowly methylated transcripts were enriched in pathways related to RNA processing, ribosome, and the proteasome (Fig. 6F). The enrichment of neurodegeneration-associated KEGG pathways in the HEK293T low-cap methylation gene set likely reflects the shared annotation of proteasome, mitochondrial, and ER stress genes across these disease pathways (see gene names in Data S6). However, because dysfunction of these processes and impaired SAM-dependent methylation could be hallmarks of neurodegeneration^8^, it will be interesting to explore if altered cap methylation of these transcripts contributes to neurodegeneration.

To determine whether cap-unmethylated mRNAs are associated with polysomes in mammalian cells, we performed polysome profiling of HEK293T cells cultured under methionine-replete or methionine-starved conditions. As expected, methionine starvation resulted in an increase in monosomes and a concomitant decrease in polysomes (Fig. 6G). We then quantified cap dinucleotides in fractions corresponding to free mRNAs and ribosomal subunits (Free), monosomes (Mono), and polysomes (Poly). Following methionine starvation, fully methylated cap m⁷Gpppm⁶Am species were detected in both monosome and polysome fractions (Fig. 6H). Notably, the unmethylated GpppA cap was also readily detected in all fractions under methionine starvation (Fig. 6I). To further assess the association of GpppA-capped mRNAs with actively translating ribosomes, we treated cells with puromycin to disrupt polysomes (Fig. 6J). Although puromycin treatment only partially reduced polysomes under our experimental conditions, both m⁷Gpppm⁶Am and GpppA cap structures showed similar reductions in the polysome fractions (Fig. 6K). Together, these findings suggest that transcript-specific mRNA cap methylation is a conserved feature of eukaryotic gene expression, with cap-unmethylated mRNAs retaining the capacity to engage with actively translating ribosomes.

## Discussion

We demonstrate for the first time that cap-unmethylated mRNAs can be prevalent in both yeast and mammalian cells, especially under conditions where methionine and SAM are limiting, setting the stage for subsequent detailed investigations of their biological functions and fates. This finding also raises several immediate questions, which we discuss below.

Surprisingly, we observed that mRNA expression and induction levels do not correlate with cap methylation levels, suggesting that transcription and cap methylation are not necessarily coupled processes. Why might cells transcribe certain mRNAs without methylating their caps? Here, we provide the first evidence that cap-unmethylated mRNAs share key features with canonical cap-methylated transcripts—they are polyadenylated, exported, and translated. Most importantly, cap-unmethylated transcripts appear important for cell growth under stress. Based on our findings, we propose several hypotheses regarding the underlying mechanisms: cap methylation status could represent an additional means of translational regulation, as cap-unmethylated mRNAs typically exhibit reduced translation efficiency compared to their methylated counterparts^41^. Cytosolic recapping and cap methylation have been reported in mammalian cells^42^. Consistent with this possibility, the lack of a growth defect upon adding a nuclear export signal (NES) to Abd1 (Fig. S6A) suggests that cytosolic cap methylation may also occur in yeast. This could allow existing cap-unmethylated transcripts to be rapidly methylated and translated in response to environmental cues. Interestingly, ribosome-related transcripts are among the least methylated (Fig. 2G), and their translation is highly correlated with cell growth. Such mRNAs may represent a class of transcripts that are “pre-synthesized and stored,” allowing for rapid expression once nutrient conditions improve, potentially conferring a growth advantage to the cell.

Intriguingly, we also identified a subset of mRNAs that retain robust TE following a shift to minimal medium (Fig. S6B, C). We focused on mRNAs with both increased expression and strong TE, reasoning that these mRNAs are more likely to represent robust cases. These findings suggest the possibility of an alternative translation initiation mechanism that may facilitate translation of unique cap-unmethylated mRNAs. Consistent with this idea, we confirmed that the translation initiation factor eIF4E^7^ binds a cap-methylated RNA oligonucleotide, but not a cap-unmethylated RNA oligonucleotide (Fig. S6D). Although we did not identify any cap-independent translation elements enriched in cap-unmethylated mRNAs, we cannot rule out the possibility that cap-independent translation contributes to the translation of GpppN-mRNAs.

By investigating several possible determinants or correlates of mRNA cap methylation, we observed that increased H3K36me3 might predict higher cap methylation. We propose two non-mutually exclusive possibilities. First, we observed an intriguing clustering of elevated H3K36me3 signals during methionine starvation, with certain chromosomal regions (such as the ends of chromosomes) forming distinct clusters of H3K36me3 enrichment (Fig. S6E). One possible explanation is that these genomic regions are also spatially clustered within the nucleus and reside in compartments with enhanced local SAM concentration. Consequently, these regions may experience elevated levels of both histone and mRNA cap methylation. In addition, the reduced m⁷GpppA–mRNA levels detected in the *H3K36A* mutant suggest that the H3K36 methylation state itself may contribute to enhanced cap methylation. Second, we found that the SAM synthetase enzymes Sam1 and Sam2 localize not only to the cytosol but also to the nucleus. Their nuclear localization was confirmed by co-labeling with the nuclear envelope marker Nic96-mCherry and was abolished by addition of a nuclear export signal (NES) (Fig. S6F, G). Moreover, co-immunoprecipitation experiments revealed interactions between Set2 and both Sam1 and Sam2 (Fig. S6H). Together, these observations raise the possibility that Set2 recruits SAM synthetases to specific chromatin regions, thereby locally enriching SAM to support both H3K36 methylation and co-transcriptional mRNA cap methylation. This possibility warrants further investigation, as the intracellular SAM/SAH ratio is likely to have a greater impact on methyltransferase activity than either metabolite alone. Although Abd1 (and mammalian RNMT) has a relatively high *K_m_* for SAM compared with some other methyltransferases^8^, its activity may be sensitive to changes in both SAM availability and SAH-mediated inhibition.

In summary, this study represents an initial effort to investigate how mRNA cap methylation can be regulated by metabolic state and other forms of stress. We report the stable existence and potential functionality of cap-unmethylated mRNAs. Our collective observations suggest that both preferentially cap-methylated mRNAs and cap-unmethylated mRNAs are important for the cell to adapt to stress. What might be the key differences between these classes of transcripts that differ by only a single methyl group in their 5’ caps? One possibility is that cap-methylated transcripts belong to the group of genes that necessitate immediate translation. They could encode proteins that prevent damage to the cell, or perhaps rate-limiting enzymes of a biosynthetic pathway. Cells can then ensure all processes important for survival under stress are prioritized, providing an additional layer of post-transcriptional regulation. For cap-unmethylated mRNAs, enforced cap methylation during metabolic stresses is apparently disadvantageous to cells. We focused on characterizing their fundamental properties by analyzing the bulk population of cap-unmethylated transcripts in this study. However, we anticipate that these mRNAs may serve diverse functional roles under different contexts. To this end, we developed transcriptome-wide methods for both yeast and mammalian cells to assess cap methylation status at the individual mRNA level. These tools will enable the identification of specific transcripts that undergo differential cap methylation across a variety of conditions beyond nutrient stress, including developmental stages, disease states, and other physiological or environmental challenges.

## Limitations of the study

This study focused on the prototrophic CEN.PK strain of the budding yeast *Saccharomyces cerevisiae*, and two cell lines, HEK293T and Neuro-2a. Future studies using additional yeast strains and mammalian cell types might further establish the generality of these findings.

The m^7^G-IP method relies on a particular antibody used by the field and may therefore be subject to non-specific binding, which we addressed using the orthogonal Cap-Tag approach we developed. m^7^G-IP also enables estimation of methylation levels using RNAs from rich-medium-grown cells as a fully methylated reference; however, some transcripts may already be partially unmethylated under these conditions, potentially affecting absolute estimates. Cap-Tag does not require this assumption but is less suitable for accurately quantifying methylation levels.

For the *in vivo* translation assay in yeast spheroplasts, we attempted to use RT-qPCR to quantify specific RNA species recovered from spheroplasts and thereby normalize for differences in RNA electroporation efficiency. However, technical limitations, particularly the inability to extensively wash fragile spheroplasts without compromising their integrity, prevented reliable interpretation of these measurements. Because all RNAs were electroporated into yeast using the same buffer and experimental procedures, we instead relied on biological replicates to account for technical variation in electroporation efficiency.

## Resource availability

### Lead Contact

Questions regarding this study, including requests for additional information or resources, should be directed to the lead contact, Benjamin Tu

### Materials Availability

Reagents generated as part of this study will be provided upon reasonable request. Depending on the nature of the request and potential commercial applications, shipping cost and/or a completed Materials Transfer Agreement may be required.

### Data and Code Availability

All data supporting the conclusions of this study are included in the main article and supplemental materials. Sequencing datasets generated in this study have been deposited in the Gene Expression Omnibus (GEO) under the following accession numbers: GSE312836, GSE313216, GSE313217, GSE313220, GSE345978, and GSE346160.

No original code was generated for this study.

Additional information needed to reanalyze the data reported in this paper can be obtained from the lead contact upon request.

## Acknowledgments.

We thank Dr. Kathleen Collins (UC Berkeley, CA) for generously providing the original and W403AF753A BoMoC enzymes for OTTR. We thank Dr. Jingxuan Chen and Dr. Jeon Lee at the DSSR help desk (UTSW) for help with bioinformatics analysis. The BioHPC computational resource at the UTSW provided computational support for this project. We thank prior lab member Dr. Kuanqing Liu for training Dr. Zheng Xing for LC-MS/MS and polysome profiling. We thank Dr. Cunqi Ye (prior Tu lab member), Dr. Jinfan Wang (UTSW) and all current Tu lab members for helpful discussions.

This work was supported by NIH R35GM136370 and HHMI to B.P.T. and R01GM139008 to N.T.I.

## Author contributions

Conceptualization: Z.X. and B.P.T.

Methodology: Z.X., B.M.S.

Investigation: Z.X., B.M.S., A.V.F., N.T.I.

Visualization: Z.X.

Funding acquisition: B.P.T, N.T.I.

Project administration: Z.X., B.P.T

Supervision: B.P.T.

Writing – original draft: Z.X.

## STAR Methods

### Key resources tables

#### Yeast strains

| Name | Genotype | References |
| --- | --- | --- |
| *Wild Type* | *MATa CEN.PK* | <sup>43</sup> |
| *TIR1* | *MATa ho:: TIR1::KanMX6* | This study |
| *ABD1-AID\** | *MATa ho:: TIR1::KanMX6 ABD1-AID\*-1XHA::HYG* | This study |
| *ABD1-AID\*cho2Δ* | *MATa ho:: TIR1::KanMX6 ABD1-AID\*-1XHA::HYG cho2:: NAT* | This study |
| *pTEF1-ABD1-HA* | *MATa NAT-P<sub>TEF1</sub> ABD1-6XHA::kanMX6* | This study |
| *dxo1Δ* | *MATa dxo1:: kanMX6* | This study |
| *rai1Δ* | *MATa rai1:: HYG* | This study |
| *rai1Δdxo1Δ* | *MATa rai1:: HYG dxo1::kanMX6* | This study |
| *pTEF1-RAI1* | *MATa HYG-P<sub>TEF1</sub> RAI1-6XHA::kanMX6* | This study |
| *pTEF1-DXO1* | *MATa HYG-P<sub>TEF1</sub> DXO1-6XHA::KanMX6* | This study |
| *pTEF1-RAI1 pTEF-DXO1* | *MATa HYG-P<sub>TEF1</sub> DXO1-6XHA:: KanMX6 NAT-P<sub>TEF1</sub> RAI1-6XHA:: ble* | This study |
| *DCP2-AID\** | *MATa ho:: TIR1::KanMX6 DCP2-AID\*-1XHA::HYG* | This study |
| *dcs1Δ* | *MATa dcs1:: kanMX6* | This study |
| *dcs2Δ* | *MATa dcs2:: kanMX6* | This study |
| *MET16-MS2 Nic96-mCherry* | *MATa ho:: MS2CP::KanMX6 MET16-MS2 NIC96-mCherry:: NAT* | This study |
| *MET16-MS2 Dhh1-mCherry* | *MATa ho:: MS2CP::KanMX6 MET16-MS2 DHH1-mCherry:: NAT* | This study |
| *jlp1Δ* | *MATa jlp1:: HYG* | This study |
| *rpn1Δ* | *MATa rpn1:: HYG* | This study |
| *gis1Δ* | *MATa gis1:: HYG* | This study |
| *tpa1Δ* | *MATa tpa1:: HYG* | This study |
| *jhd2Δ* | *MATa jhd2:: HYG* | This study |
| *jhd1Δ* | *MATa jhd1:: HYG* | This study |
| *ecm5Δ* | *MATa ecm5:: HYG* | This study |
| *Pab1-GFP* | *MATa PAB1-GFP:: KanMX6* | This study |
| *Pub1-GFP* | *MATa PUB1-GFP:: KanMX6* | This study |
| *Ded1-GFP* | *MATa DED1-GFP:: KanMX6* | This study |
| *Dcp2-GFP* | *MATa DCP2-GFP:: KanMX6* | This study |
| *Dhh1-GFP* | *MATa DHH1-GFP:: KanMX6* | This study |
| *Edc3-GFP* | *MATa EDC3-GFP:: KanMX6* | This study |
| *BET1-MS2 Dhh1-mCherry* | *MATa ho:: MS2CP::KanMX6 BET1-MS2 DHH1-mCherry:: NAT* | This study |
| *RPN12-MS2 Dhh1-mCherry* | *MATa ho:: MS2CP::KanMX6 RPN12-MS2 DHH1-mCherry:: NAT* | This study |
| *ABD1-HA* | *MATa ABD1-6XHA:: KanMX6* | This study |
| *HO-GEV-NSP14-HA* | *MATa ho::GEV-NSP14-6xHA::NAT* | This study |
| *HO-GEV-NSP14-GFP* | *MATa ho::GEV-NSP14-GFP::NAT* | This study |
| *set2Δ* | *MATa set2:: KanMX6* | 25 |
| *H3K36A* | *HHT1-HHF1-H3K36A::hygMX, HHT2-HHF2-H3K36A::natMX* | 25 |
| *HOG1-HA* | *MATa HOG1-6XHA:: KanMX6* | This study |
| *SLT2-HA* | *MATa SLT2-6XHA:: KanMX6* | This study |
| *HOG1-AID\** | *MATa ho:: TIR1::KanMX6 HOG1-AID\*-1XHA::HYG* | This study |
| *SLT2-AID\** | *MATa ho:: TIR1::KanMX6 SLT2-AID\*-1XHA::HYG* | This study |
| *HOG1-FLAG* | *MATa HOG1-3XFLAG:: KanMX6* | This study |
| *SLT2-FLAG* | *MATa SLT2-3XFLAG:: KanMX6* | This study |
| *TIR1cho2Δ* | *MATa ho:: TIR1::KanMX6 c2:: NAT* | This study |
| *bem2Δ* | *MATa bem2::HYG* | This study |
| *cla4Δ* | *MATa cla4::HYG* | This study |
| *mid2Δ* | *MATa mid2::HYG* | This study |
| *msb2Δ* | *MATa msb2::HYG* | This study |
| *sac7Δ* | *MATa sac7::HYG* | This study |
| *ssk2Δ* | *MATa ssk2::HYG* | This study |
| *yck2Δ* | *MATa yck2::HYG* | This study |
| *TIR1 BEM2-FLAG* | *MATa ho:: TIR1::KanMX6 BEM2-3XFLAG:: NAT* | This study |
| *TIR1 CLA4-FLAG* | *MATa ho:: TIR1::KanMX6 CLA4-3XFLAG:: NAT* | This study |
| *TIR1 MID2-FLAG* | *MATa ho:: TIR1::KanMX6 MID2-3XFLAG:: NAT* | This study |
| *TIR1 MSB2-FLAG* | *MATa ho:: TIR1::KanMX6 MSB2-3XFLAG:: NAT* | This study |
| *TIR1 SAC7-FLAG* | *MATa ho:: TIR1::KanMX6 SAC7-3XFLAG:: NAT* | This study |
| *TIR1 SSK2-FLAG* | *MATa ho:: TIR1::KanMX6 SSK2-3XFLAG:: NAT* | This study |
| *TIR1 YCK2-FLAG* | *MATa ho:: TIR1::KanMX6 YCK2-3XFLAG:: NAT* | This study |
| *TIR1 MSS4-FLAG* | *MATa ho:: TIR1::KanMX6 MSS4-3XFLAG:: NAT* | This study |
| *TIR1 RSP5-FLAG* | *MATa ho:: TIR1::KanMX6 RSP5-3XFLAG:: NAT* | This study |
| *TIR1 PKC1-FLAG* | *MATa ho:: TIR1::KanMX6 PKC1-3XFLAG:: NAT* | This study |
| *ABD1-AID\* BEM2-FLAG* | *MATa ho:: TIR1::KanMX6 ABD1-AID\*-1XHA::HYG BEM2-3XFLAG:: NAT* | This study |
| *ABD1-AID\* CLA4-FLAG* | *MATa ho:: TIR1::KanMX6 ABD1-AID\*-1XHA::HYG CLA4-3XFLAG:: NAT* | This study |
| *ABD1-AID\* MID2-FLAG* | *MATa ho:: TIR1::KanMX6 ABD1-AID\*-1XHA::HYG MID2-3XFLAG:: NAT* | This study |
| *ABD1-AID\* MSB2-FLAG* | *MATa ho:: TIR1::KanMX6 ABD1-AID\*-1XHA::HYG MSB2-3XFLAG:: NAT* | This study |
| *ABD1-AID\* SAC7-FLAG* | *MATa ho:: TIR1::KanMX6 ABD1-AID\*-1XHA::HYG SAC7-3XFLAG:: NAT* | This study |
| *ABD1-AID\* SSK2-FLAG* | *MATa ho:: TIR1::KanMX6 ABD1-AID\*-1XHA::HYG SSK2-3XFLAG:: NAT* | This study |
| *ABD1-AID\* YCK2-FLAG* | *MATa ho:: TIR1::KanMX6 ABD1-AID\*-1XHA::HYG YCK2-3XFLAG:: NAT* | This study |
| *ABD1-AID\* MSS4-FLAG* | *MATa ho:: TIR1::KanMX6 ABD1-AID\*-1XHA::HYG MSS4-3XFLAG:: NAT* | This study |
| *ABD1-AID\* RSP5-FLAG* | *MATa ho:: TIR1::KanMX6 ABD1-AID\*-1XHA::HYG RSP5-3XFLAG:: NAT* | This study |
| *ABD1-AID\* PKC1-FLAG* | *MATa ho:: TIR1::KanMX6 ABD1-AID\*-1XHA::HYG PKC1-3XFLAG:: NAT* | This study |
| *RAI1-HA* | *MATa RAI1-6XHA:: KanMX6* | This study |
| *DXO1-HA* | *MATa DXO1-6XHA:: KanMX6* | This study |
| *Set2-FLAG Sam1-HA* | *MATa SET2-3XFLAG:: HYG SAM1-6XHA:: KanMX6* | This study |
| Set2-FLAG Sam2-HA | *MATa SET2-3XFLAG:: HYG SAM2-6XHA:: KanMX6* | This study |
| Sam1-HA | *MATa SAM1-6XHA:: KanMX6* | This study |
| Sam2-HA | *MATa SAM2-6XHA:: KanMX6* | This study |
| SAM1-eGFP<br>Nic96-mCherry | *MATa SAM1-eGFP:: KanMX6 NIC96-mCherry::HYG* | This study |
| SAM2-eGFP<br>Nic96-mCherry | *MATa SAM2-eGFP:: KanMX6 NIC96-mCherry::HYG* | This study |
| SAM1-NES-eGFP<br>Nic96-mCherry | *MATa SAM1-NES-eGFP:: KanMX6 NIC96-mCherry::HYG* | This study |
| SAM2-NES-eGFP<br>Nic96-mCherry | *MATa SAM2-NES-eGFP:: KanMX6 NIC96-mCherry::HYG* | This study |
| Cdc33-HA | *MATa CDC33-6XHA:: KanMX6* | This study |

#### Plasmids

| Name | Description | References |
| --- | --- | --- |
| pFA6a-Hyg | To knock out a gene of interest (GOI) with the Hyg gene | 44 |
| pFA6a-KanMX6 | To knock out a GOI with the KanMX6 gene | 44 |
| pFA6a-HA-KanMX6 | To tag a GOI with 6XHA tag, kanamycin resistance | This study |
| pFA6a-HA-Hyg | To tag a GOI with 6XHA tag, hygromycin resistance | This study |
| pFA6a-HA-ble | To tag a GOI with 6XHA tag, phleomycin resistance | This study |
| pFA6a-FLAG-Nat | To tag a GOI with 3XFLAG tag, nourseothricin resistance | This study |
| pFA6a-TEF1-OM-KanMX6 | To insert the TEF1 promoter in front of the CDS of a GOI, kanamycin resistance | This study |
| pFA6a-TEF1-OM-Nat | To insert the TEF1 promoter in front of the CDS of a GOI, nourseothricin resistance | This study |
| HO-NSP14-6XHA-Nat-HO | To insert NSP14-6XHA at the HO locus | This study |
| HO-NSP14-yeGFP-Nat-HO | To insert NSP14-yeGFP at the HO locus | This study |
| pFA6a-yeGFP-KanMX6 | To tag a GOI with yeGFP tag | This study |
| pFA6a-NES-yeGFP-KanMX6 | To tag a GOI with NES-yeGFP tag | This study |
| pNHK53 | Template for osTIR1 PCR | 24 |
| pHyg-AID* | Template for AID* PCR | 24 |
| HO-OsTIR1-KanMX6-HO | To insert OsTIR1 at the HO locus | This study |
| pHyg-AID*-HA | To tag a gene of interest with AID*-1XHA tag | This study |
| pET296-YcpLac111 CYC1p-MCP-NLS-2xyeGFP | Template for CYC1p-MCP-NLS-2xyeGFP PCR | 33 |
| HO-CYC1p-MCP-NLS-yeGFP-Hyg-HO | To insert CYC1p-MCP-NLS-yeGFP at the HO locus | This study |
| pET259-pUC57 24xMS2V6 | To tag a gene of interest with 24XMS2V6 | 33 |
| pSH65 | To induce Cre recombinase expression under GAL promoter | 33 |
| pACT-GEV | To PCR the ACT-GEV fragment for integration | 45 |
| IVT-MET16-2XFLAG | To PCR MET16 UTRs and CDS for in vitro transcription | This study |
| IVT-MET16UTR-NanoLuc | To PCR MET16 UTRs and NanoLuc CDS for in vitro transcription | This study |

#### FISH probes

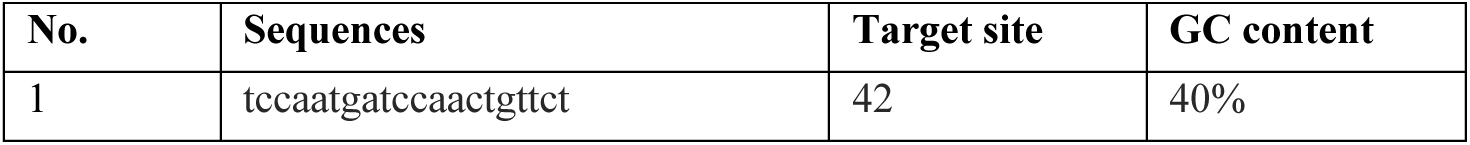
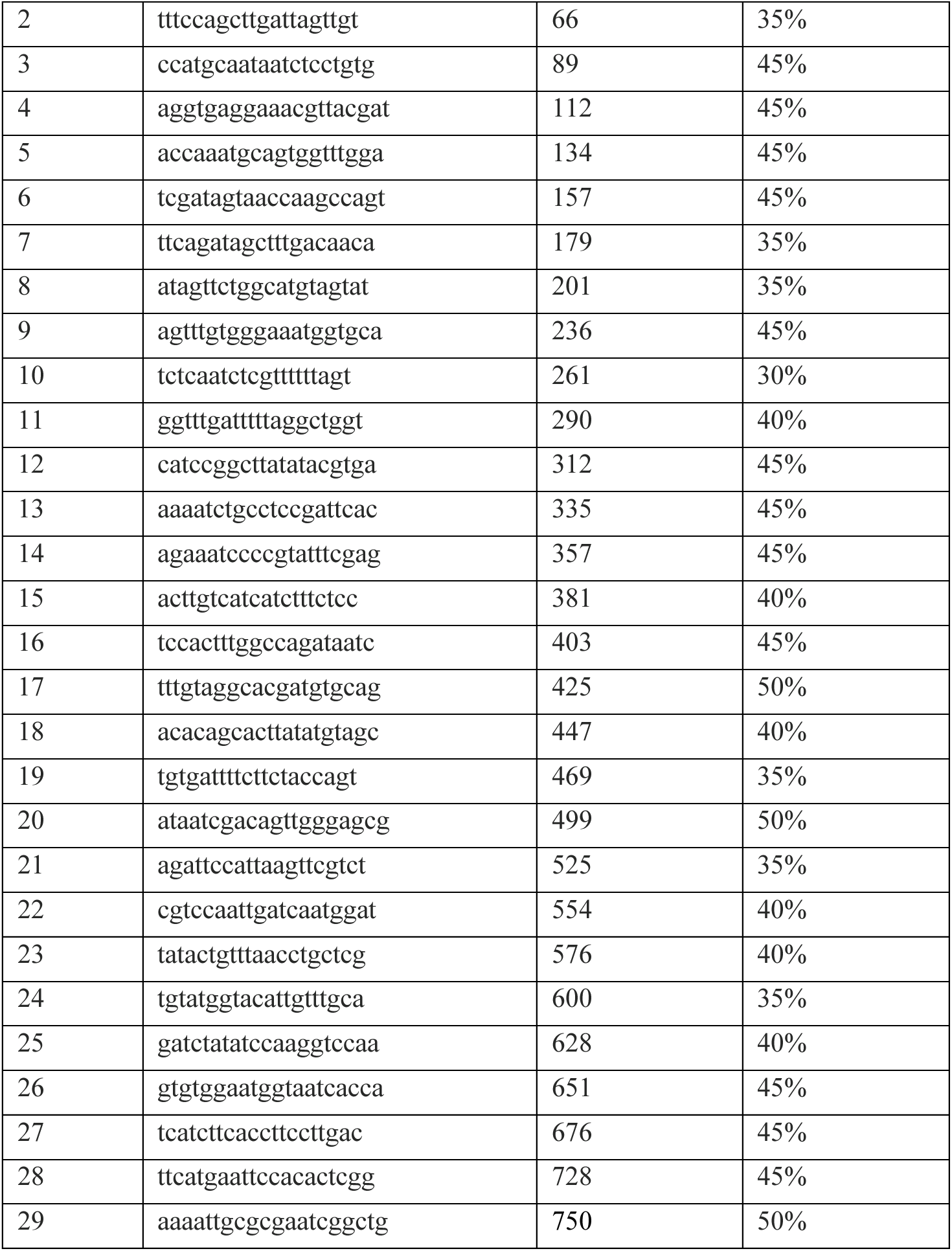

#### Antibodies

1 Antibodies
| Target | Source |
| --- | --- |
| m<sup>7</sup>G | Sigma, NA02 |
| HA | Cell Signaling Technology, 3724S |
| FLAG | Sigma, F3165 |
| p-Hog1 | Cell Signaling Technology, 9211S |
| p-Slt2 | Cell Signaling Technology, 4370S |
| Total Hog1 | Santa Cruz Biotechnology, sc-165978 |
| Total Slt2 | Santa Cruz Biotechnology, sc-133189 |
| GFP | Roche, 11814460001 |
| Rpn10 | Abcam, ab98843 |
| H3K4me3 | Abcam, ab8580 |
| H3K36me3 | Abcam, ab9050 |
| H3K79me3 | Abcam, ab2621 |
| H3 | Sigma, 05-928 |
| Rpb1 | Biolegend, 664906 |

### Method details

#### Yeast growth

Yeast cultures were grown at 30°C with 300 rpm shaking. Cultures were typically inoculated the day before, diluted to OD_600_ ∼ 0.2 the next morning and allowed to grow to log phase (OD_600_ ∼ 0.8) for various experiments. YPL is a rich medium containing 10 g yeast extract (BioBasic, G0961) and 20 g peptone (Bacto, 211677) in 1 L, supplemented with 2% lactate. SL is a minimal medium lacking amino acids and containing 6.7 g yeast nitrogen base (BD, 291920) in 1 L, with 2% lactate. For media switch, cells were grown in YPL, part of the culture was harvested by centrifugation and flash frozen in liquid nitrogen when applicable, another part was washed once with SL, and resuspended in the same volume of SL. Heavy guanine (Cambridge Isotope Laboratories, CNLM-3990-25), thiolutin (Cayman, 11350), and H_2_O_2_ (Sigma, 216763) were also used when indicated. For serial dilution growth assays, yeast cells were washed with sterile water and spotted on indicated agar plates in 6-fold dilutions, starting with OD_600_ = 0.1. For growth curve analysis, yeast cells were washed with desired growth media and then seeded 100 µL/well in flat bottom 96-well sterile plates (CELLTREAT, 229596). OD_600_ was measured using a Tecan Spark plate reader at 30 min intervals with continuous shaking and a magnetic lid lifter for better measurement and aeration.

#### Cap LC-MS/MS

Total RNA was extracted from yeast using the acid phenol method. When applicable, poly (A)+ RNA was purified from total RNA using Dynabeads mRNA purification kit (Invitrogen, 61006). 50 µg of total RNA or 1 µg of poly (A)+ RNA was digested in 110 µL reaction containing 7 µL acidic buffer (0.1 M NaOAc pH 6.8, 20 mM ZnCl_2_), 1 µL RNase A (5 µg/µL, Biosearch, MRNA092) and 1 µL Nuclease P1 (1.5 U/µL, Sigma, N8630). After incubation at 37°C for 4 h, 7 µL basic buffer (0.3 M NaOAc pH 7.8) and 1 µL calf intestinal phosphatase (NEB, M0525) was added and the reaction was allowed to continue at room temperature overnight. The reaction was filtered using Amicon ultra 0.5 ml filtration units (10K MWCO). The flow-through, containing the digested nucleosides and cap structures, was collected, and 50 µL was injected into a Synergi 4 µm Fusion-RP 80A 150 X 2 mm column (Phenomenex, 00F-4424-B0) using a Shimadzu high performance liquid chromatography (HPLC) system. HPLC was operated at 0.5 mL/min with 10 mM tributylamine pH 5 (buffer A) and 100% methanol (buffer B). Gradient elution was performed as follows: 0.01 min, 0% B, 11 min, 50% B, 13 min, 100% B, 21 min, 100% B, 22 min, 0% B, 30 min, 0% B. Analytes were detected using SCIEX 5500 QTRAP mass spectrometer with negative mode and multiple reaction monitoring (MRM) using transitions listed below. Analytes were quantified using the SCIEX OS software.

#### Yeast metabolite analysis

Metabolites were extracted and dried as described^46^. Dried metabolites were resuspended in buffer A (5 mM NH_4_OAc pH 5.5 in water) and filtered with a 0.2 µm PVDF syringe filter before injecting into a Synergi 4 µm Fusion-RP 80A 150 X 2 mm column. HPLC was operated at 0.5 mL/min with buffer A and buffer B (5 mM NH_4_OAc in methanol). The following gradient elution was performed: 0.01 min, 0% B, 5 min, 0% B, 6 min, 1% B, 7 min, 3% B, 8 min, 5% B, 14 min, 25% B, 16 min, 50 % B, 18 min, 100% B, 22 min, 100% B, 23 min, 0% B, 28 min, 0% B. Metabolites were detected with 5500 QTrap with positive mode and MRM using transitions listed below, and quantified with SCIEX OS.

#### Mass spectrometry transitions for cap and metabolites analysis

| Analyte | Mode | Q1 | Q3 |
| --- | --- | --- | --- |
| m<sup>7</sup>GpppA | - | 785 | 408 |
| GpppA | - | 771 | 408 |
| m<sup>7</sup>GpppG | - | 801 | 424 |
| GpppG | - | 787 | 424 |
| m<sup>7</sup>Gpppm<sup>6</sup>A<sub>m</sub> | - | 813 | 436 |
| m<sup>7</sup>GpppAm | - | 799 | 422 |
| GpppAm<sup>6</sup>A<sub>m</sub> | - | 799 | 436 |
| GpppAm | - | 785 | 422 |
| m<sup>7</sup>GpppG<sub>m</sub> | - | 815 | 438 |
| GpppG<sub>m</sub> | - | 801 | 438 |
| adenosine | - | 266 | 134 |
| guanosine | - | 282 | 150 |
| C13 m<sup>7</sup>GpppA | - | 788 | 620 |
| C13 m<sup>7</sup>GpppA | - | 788 | 408 |
| C13 GpppA | - | 774 | 408 |
| C13 GpppA | - | 774 | 427 |
| C13 guanosine | - | 285 | 153 |
| Propargyl-GpppA | - | 809 | 408 |
| SAM | + | 399 | 250 |
| SAH | + | 385 | 136 |

#### Whole cell lysate preparation and Western blot

Yeast cells were harvested by centrifugation and resuspended in 20% trichloroacetic acid (TCA) and incubated on ice for 15 min and washed once in cold acetone. TCA-treated cell pellet was lysed by bead-beating (3 X 30 s) in lysis buffer (50 mM Tris-HCl pH 7.5, 6 M urea, 1% SDS, 5 mM EDTA, 1 mM DTT, 1 mM PMSF, 10 µM leupeptin, 5 µM pepstatin A, and 1X protease inhibitor cocktail (Roche, 11697498001)). For phosphorylation western blots, 20 mM NaF and 1 mM NaNO_3_ were added. After bead-beating, lysate was incubated at 75°C for 5 min. Mammalian cells were washed once with 1× PBS, scraped using a cell scraper, and collected by centrifugation at 500 × g. Cell pellets were lysed in lysis buffer (Cell Signaling Technology, #9083) supplemented with Protease/Phosphatase Inhibitor Cocktail (Cell Signaling Technology, #5872). Lysates were clarified by centrifugation at 14,000 × g, and the supernatants were collected for analysis. Western blotting was performed using standard procedures.

#### Dot-blot analysis

Total poly (A)+ RNAs were isolated as described above. mRNAs were diluted to desired concentrations, heated at 70°C for 2 min, snap-cooled on ice, and 2 µL of mRNA solutions were dropped directly onto Hybond-N+ membrane (Cytiva, RPN303B) and allowed to dry briefly. The membrane was crosslinked in a Stratalinker 200 UV crosslinker twice using the auto-crosslink mode (1200 microjoules [x100]). For blotting, the membrane was washed in wash buffer (1x PBS, 0.02% Tween-20) for 5 min at room temperature, incubated with blotting buffer (1x PBS, 0.02% Tween-20, 2% BSA) for 1 h at room temperature, incubated with anti-m^7^G clone K121 antibody (1:35) overnight at 4°C, washed 3X for 5 min each in wash buffer at room temperature, incubated with goat anti-mouse IgG-HRP (1:10000) for 1 h at room temperature, and washed 4X for 10 min each with wash buffer before imaging using standard Western protocols. To visualize total mRNA on the membrane, a methylene blue solution (0.2 % methylene blue, 0.4 M NaOAc, 0.4 M acetic acid) was incubated with the membrane for 30 min at room temperature and then washed with MilliQ water until desired signal/background reached.

#### m^7^G immunoprecipitation

Total yeast RNA was extracted, treated with Turbo DNase (Ambion, AM2238) and extracted with acid phenol again. Poly(A)+ RNA was enriched as described above. 50 µl of Dynabeads protein G (ThermoFisher, 10003D) was washed 2X with RNA immunoprecipitation (RIP) buffer (25 mM Tris-HCl pH 7.5, 200 mM NaCl, 5 mM EDTA, 0.1% Tween-20, 0.5 mM DTT). To pre-bind antibody, washed beads were mixed with 10 µg m^7^G K121 antibody (Sigma, NA02) or 10 µg mouse IgG in 200 µL of RIP buffer. The mixture was incubated at 4°C for 2 h with rotation, and washed 2X with 500 µL RIP buffer. 300 ng poly(A)+ RNA was added to the beads in 200 µL RIP buffer with 0.1U/µL SUPERase·In (ThermoFisher, AM2696), and incubated at 4°C overnight with rotation. The next day, the beads were washed 3X with 1 mL RIP buffer. Finally, the washed immunoprecipitates were subjected to acid phenol extraction, and precipitated with ethanol and glycogen. The RNA pellet was resuspended in 20 µL water and two biological replicates were subjected to strand-specific RNA-sequencing with Novogene. Genes with fewer than 10 reads in any sample were excluded from further analysis. Data were normalized to the total counts of ERCC RNA spike-ins (ThermoFisher, 4456740). Percent methylation of RNAs from rich medium samples was arbitrarily set to 100% (i.e., (IP^rich^/input^rich^)*X=% methylation^rich^=100%), and % methylation of RNAs from minimal medium samples was calculated using the same scaling factor: % methylation^min^=(IP^min^/input^min^)*X. Because the % methylation in rich medium was arbitrarily set, this calculation yielded 229 transcripts in minimal medium with values exceeding 100%. For these transcripts, the % methylation was capped at 100%. For the *pTEF1-ABD1-HA* strain, m^7^G-IP was performed as described above. Data were normalized by library size. During % methylation calculation, 626 transcripts with calculated values exceeding 100% were capped at 100%.

#### In vitro Cap-Tag

Poly(A)+ RNA was extracted from yeast cells as described above. RNA was DNase I-treated and purified with RNA Clean & Concentrator (Zymo Research, R1017). 500 ng poly(A)+ RNA was labeled by incubating at 37°C for 35 min with 0.05 mM propargylic *Se*-adenosyl-*L*-selenomethionine (ProSeAM)^28^ (WuXi AppTec) and 1 µL Vaccinia capping enzyme (NEB, M2080S) (omitted in the background control group) in a 50 µL reaction volume with buffer containing 50 mM Tris-HCl pH 7.5, 5 mM KCl, and 1 mM DTT. Labeled RNA was cleaned with the Zymo RNA Clean & Concentrator kit and eluted in 10 µL water. Click-chemistry reagents were obtained from Jena Bioscience (CLK-071) and reaction was performed per manufacturer instructions. Briefly, 2 µL CuSO_4_ (100mM) and 20 µL BTTAA (50 mM) were mixed and incubated at room temperature for 5 min. 10 µL labeled RNA, 57.5 µL 100 mM Na-Phosphate buffer pH 7, and 0.5 µL 1 mM Picolyl-Azide-PEG4-Biotin were mixed, and then added to the first part of reaction. 100 mM NaAscorbate was added to start the reaction, and then incubated at 37°C for 35 min. Samples were cleaned with the RNA Clean & Concentrator kit and eluted in 17 µL water. 1 µL of RNA was saved at this step as input.

For each pull-down, 10 µL Dynabeads M-280 Streptavidin (Invitrogen,11205D) was washed 3X with bind buffer (50 mM Tris-HCl pH 7.5, 1 M NaCl, 5 mM EDTA, 0.2% Triton X-100), blocked for 2 h with blocking buffer (10 mM Tris-HCl pH 7.5, 0.2 mg/mL BSA, 0.2 mg/mL yeast tRNA, 0.01 mg/mL glycogen) at room temperature, and washed 3X again with bind buffer. 15 µL sample from the Click reaction was incubated with the beads in 100 µL bind buffer for 45 min at room temperature. The beads were washed 3X with bind buffer at room temperature, 1X with bind buffer with 4 M NaCl at 42°C for 3 min, and 1X with 1 X PBS. The washed beads were then eluted with 50 µL elution buffer (95% formamide, 10 mM EDTA) at 65°C for 5 min. The beads together with the eluate were purified with TRIzol according to manufacturer instructions and two biological replicates were subjected to RNA sequencing with Novogene. After read counting, genes with fewer than 10 reads in any sample were excluded from further analysis. Percent input was calculated using the formula: % input = (count_pulldown / count_input / 22.5) × 100. 22.5 was the dilution factor for input samples. Background levels were estimated using both spiked-in CleanCap eGFP mRNA (TriLink, L-7601) and RNA-seq data from ΔMTase samples.

For mammalian cells, Cap-Tag was performed with the following modifications: (1) ProSeAM was used at a final concentration of 5 µM in the capping reaction; (2) THPTA (Jena Bioscience, CLK-1010-25) was used as the ligand for the click reaction instead of BTTAA due to longer shelf life when stored at -20 °C; and (3) two methylated (TriLink, L-7601, L-7203) and two unmethylated spike-in RNAs (*in vitro* transcribed) were added at 0.5 ng each per reaction. Raw read counts were normalized using ERCC spike-in controls, and genes with fewer than 10 reads in any sample were excluded from further analysis. Percent input (% input) was then calculated. Cap methylation status was determined relative to the methylated spike-in controls. Specifically, the mean ERCC-normalized % input of the two methylated spike-ins was used as the reference (mean = 5.57%, SD = 0.49%). Thresholds were defined as follows: low methylation, genes with a mean % input > mean + 5 SD of the methylated spike-in reference (i.e., >8.02%; *n* = 1,219 genes); and high methylation, genes with a mean % input < mean − 5 SD of the methylated spike-in reference (i.e., <3.12%; *n* = 5,434 genes).

#### RNA-sequencing data analysis

Data analysis for yeast datasets was done by Novogene. Briefly, raw data in fastq format was processed and clean reads were obtained by removing reads containing adapter, reads containing poly-N and low-quality reads. Reference genome *Saccharomyces cerevisiae* (R64-1-1) and gene model annotation files were downloaded from ensembl. Index of the reference genome was built, and paired-end clean reads were aligned to the reference genome using Hisat2 v2.0.5. FeatureCounts v1.5.0-p3 was used to count the number of reads mapped to each gene.

For human datasets, raw reads were processed using the Galaxy platform (usegalaxy.org). Reads were aligned to the human reference genome (GRCh38) using STAR (v2.7) in 2-pass mode. The following STAR parameters were used: sjdbOverhang = 149 (read length − 1), chimeric junction detection disabled, default mismatch tolerance. Gene-level read counts were quantified using FeatureCounts with the GRCh38 Ensembl GTF annotation.

#### ChIP-seq and data analysis

100 OD₆₀₀ units of cells were crosslinked in 1% formaldehyde for 15 min and quenched with 125 mM glycine for 10 min at room temperature. Cell pellets were washed twice with buffer containing 100 mM NaCl, 10 mM Tris-HCl (pH 8.0), 1 mM PMSF, and 1 mM benzamidine-HCl, then snap-frozen in liquid nitrogen. Frozen pellets were resuspended in 0.45 ml ChIP lysis buffer (50 mM HEPES-KOH pH 7.5, 500 mM NaCl, 1 mM EDTA, 1% Triton X-100, 0.1% sodium deoxycholate, 0.1% SDS, 1 mM PMSF, 10 μM leupeptin, 5 μM pepstatin A, and protease inhibitor cocktail (Roche)) and lysed by bead beating. For histone modifications, lysates were sonicated for 16 cycles (30 s on, 1 min off, high output) using a Bioruptor (Diagenode). For Abd1-HA and Rpb1 ChIP samples, sonication was performed with a Pixul system (Active Motif) at 50 pulses, 1 kHz pulse-rate frequency, and 20 Hz burst rate for 40 min. Sonicated lysates were clarified by two sequential centrifugations at 15,000 × g for 10 min each. For Abd1-HA and Rpb1 ChIP, 240 µL of lysate was incubated overnight with 10 µg anti-HA (Cell signaling, 3724S) or 1.5 µg anti-Rpb1 antibody (Biolegend, 664906), respectively. For histone ChIP, 50 µL of lysate was incubated overnight with 2 µg of antibody (Abcam, ab8580, ab9050, and ab2621, Sigma, 05-928). Immunoprecipitated complexes were captured with 25 µL magnetic beads for 1.5 h and washed twice with ChIP lysis buffer, once with 1 mL deoxycholate buffer (10 mM Tris-HCl, 0.25 M LiCl, 0.5% deoxycholate, 1 mM EDTA), and once with 1 mL TE buffer (10 mM Tris-HCl pH 8.0, 1 mM EDTA). Chromatin was eluted with TES buffer (10 mM Tris-HCl pH 8.0, 1 mM EDTA, 1% SDS), and crosslinks were reversed at 65 °C overnight. Eluates were treated with proteinase K (1.25 mg/mL) and RNase A (0.4 mg/mL) for 2 h at 37 °C, and DNA was purified using a QIAGEN PCR purification kit. Two biological replicates for each experiment were sent to Novogene for library construction and sequencing. ChIP-seq analyses were performed on the BioHPC cluster at UT Southwestern Medical Center. Raw sequencing data were assessed for quality using FastQC (v0.11.9), and adapters and low-quality reads were trimmed with Trim Galore! (v0.6.7). Reads from the Abd1-HA and Rpb1 datasets were aligned to a composite *S. cerevisiae* (sacCer3) and *D. melanogaster* (dm6) reference genome using BWA-MEM2 to enable spike-in normalization. Reads from histone methylation (H3K4me3, H3K36me3, H3K79me3) and total H3 datasets were aligned to the sacCer3. PCR duplicates were removed using the MarkDuplicates tool from Picard (v2.26.10) prior to downstream analyses. Significantly enriched regions were identified with MACS3 (v3.0.0). Peaks for Abd1-HA and Pol II (Rpb1) were called against their respective input DNA controls, whereas histone modification peaks were called against the total H3 control to account for nucleosome occupancy. Signal tracks for visualization were generated using deepTools (v3.5.1). Abd1-HA binding relative to Pol II was quantified as the total Abd1-HA signal across its called peaks divided by the normalized Pol II signal across the gene body. Differential histone modification analysis was performed using DiffBind with a consensus peak set derived from MACS3 outputs. Read counts within consensus peaks were quantified per replicate, and differential enrichment was assessed with the integrated DESeq2 method. Sites with a false discovery rate (FDR) < 0.05 were considered differentially bound.

#### Fluorescence *in situ* hybridization (FISH)

RNA FISH was performed using Stellaris RNA FISH probes with CAL Fluor Red 610 Dye (LGC Biosearch Technologies, set of 29 probes) according to the manufacturer’s protocol with minor modifications. Cells were cultured in appropriate media to OD_600_ of 0.4. Fixation was performed by adding 5 mL of 37% formaldehyde (Electron microscopy sciences, 15686) directly to 45 mL of culture and incubating for 45 min at room temperature. Cells were pelleted at 1600 x g for 4 min, washed once with ice-cold fixation buffer (1.2 M sorbitol, 0.1 M K_2_HPO_4_, pH 7.5), and resuspended in 1 mL fixation buffer containing 10 µl Zymolyase 100T. Digestion proceeded at 30°C for around 45 min until >80% of cells appeared phase dark. Digested cells were pelleted at 400 x g for 5 min and washed twice with ice-cold fixation buffer. Permeabilization was performed by resuspension in 70% ethanol and storage at 4 °C overnight. Prior to hybridization, cells were pelleted and resuspended in hybridization buffer (10% formamide in Stellaris hybridization buffer) containing 125 nM Stellaris probe set targeting the RNA of interest (*MET16*). Hybridization was carried out at 30°C overnight in the dark. Following hybridization, cells were washed with wash buffer A (10% formamide in 1X wash buffer A) for 30 min at 30°C, stained with 5 ng/mL DAPI (ThermoFisher, 62248) in wash buffer A for 30 min at 30°C, and rinsed briefly in wash buffer B. Cells were resuspended in Vectashield mounting medium and imaged immediately.

#### Microscopy

Images were acquired using a Zeiss Axio Observer.Z1 inverted confocal fluorescence microscope with a Plan-Apochromat 100×/1.40 oil DIC M27 objective with immersion oil. For live-cell imaging, yeast cells were grown in appropriate medium and imaged immediately at room temperature. Cells grown in YPL were washed in synthetic complete medium supplemented with lactate (SCL) prior to imaging, while cells grown in SL were imaged directly. Imaging was performed with mCherry channel (excitation 587 nm, emission 610 nm; 543–658 nm detection), and eGFP channel (excitation 488 nm, emission 509 nm; 481–543 nm detection) and DAPI channel. For RNA FISH, CAL Fluor Red 610 Dye was detected using the mCherry channel. Images were acquired with confocal pinhole set to 93 µm. Data were acquired and exported using ZEN Blue software. The same brightness and contrast were applied across all images.

#### Polysome profiling

Polysome profiling was performed as described^39^. Briefly, cells were grown at indicated conditions and cycloheximide (CHX) was added to 0.1 mg/mL at the time of harvest. Cells were rapidly chilled and incubated on ice for 5 min before centrifugation at 4000 x g for 2 min. Cell pellets were frozen in liquid nitrogen and stored in -80°C. Cell pellets were washed 2X with polysome extraction buffer (PEB = 10 mM Tris-HCl pH 7.5, 140 mM KCl, 5 mM MgCl_2_, 1% Triton X-100, 0.5 mM DTT, and 0.1 mg/mL CHX). Washed cells were lysed in PEB by bead beating method (3X, 30 s per round with 2 min on ice in between). Lysate was centrifuged at 8000 x g for 5 min at 4°C and supernatant was collected. A 10%–50% (w/v) sucrose gradient was prepared using BIOCOMP Gradient Station. Cleared lysates (normalized using A_260_ among samples) were loaded on top of the sucrose gradient and centrifuged in a 12 mL centrifuge tube (Seton, 7030) at 41,000 x g for 2 h at 4°C. Polysome fractions (0.7 mL each) were collected and A260 was measured using BIOCOMP Gradient Station. For RNA extraction, individual or pooled fractions were precipitated with ethanol and then briefly dissolved in 50 µL water with 100 U/µL rRNasin® (Promega, N2511), followed by RNA isolation with TRIzol reagent per manufacturer instructions.

#### Northern blot following polysome profiling

Northern blotting was performed as previously described^47^. RNA from each fraction was resuspended in equal volume and denatured in formaldehyde loading solution (500 µL formamide, 176 µL 37% formaldehyde, and 50 µL 20× MOPS) at 65–70 °C for 10 min and chilled on ice for 5 min. RNA samples were resolved on a 1.2% denaturing formaldehyde-agarose gel (1.8 g agarose, 7.5 ml 20× MOPS, and 8.04 mL 37% formaldehyde in 150 mL total volume) ran at 4 V/cm for 3 h. After electrophoresis, the gel was soaked in 50 mM NaOH for 20 min and rinsed thoroughly with MilliQ water. Residual formaldehyde was removed by three sequential 5 min rinses in 2× SSC (30 mM sodium citrate pH 7.0 and 300 mM NaCl). RNA was transferred overnight to a Hybond-N+ nylon membrane (Cytiva, RPN303B) via capillary blotting in 10×SSC. After transfer, RNA was UV cross-linked to the membrane (0.12 J, 2X) using a UV crosslinker. Membranes were stained with 0.02% methylene blue in 0.3 M sodium acetate (pH 5.0) to visualize rRNA. For hybridization, membranes were pre-incubated at 65 °C for 1 h in hybridization buffer (50% formamide, 5× SSPE (50 mM NaH_2_PO_4_ pH 7.4, 750 mM NaCl, 5 mM EDTA, 0.125% SDS), 1× Denhardt’s solution (0.2 g/L Ficoll 400, 0.2 g/L polyvinylpyrrolidone, 0.2 g/L BSA). Biotinylated DNA probe (for *MET16* mRNA: 5′-Biotin-GGACGTTCGAGCAGGTTAAACAGTATATAGATGCAAACAA-3′) was added at 10 nM and hybridized at 65 °C for 1 h followed by overnight incubation at 37 °C. Membranes were washed twice with 2× SSC and 0.1% SDS at room temperature and twice with 0.2× SSC and 0.1% SDS at 37 °C for 20 min each. Detection was performed using the Chemiluminescent Nucleic Acid Detection Module (Thermo Fisher, 89880) following the manufacturer’s protocol.

#### Ribosome profiling

300 mL of yeast culture were grown in YPL to OD 600 ∼ 0.8 and harvested, or switched to SL for 1 h and harvested. At time of harvesting, cells were poured into a pre-warmed filtration apparatus and filtered through a 0.45 µm cellulose nitrate membrane and immediately scraped into a 50 mL conical tube containing liquid nitrogen. Frozen cells were combined with polysome lysis buffer without cycloheximide, dripped into the liquid nitrogen, and then subjected to cryogrinding, as described^48^. The resulting pulverized cells were thawed on ice and clarified by centrifugation at 20,000xg for 30 min at 4°C. The supernatant was transferred to new tubes and frozen in liquid nitrogen before long-term storage at −80°C. Ribosome profiling was performed as described^49^, with modifications. Briefly, total RNA was quantified with the Quant-iT RiboGreen RNA kit (R11490, ThermoFisher) relative to the rRNA reference standard provided with the kit and fluorescence was measured by GloMax-Multi Jr Detection System (E6070, Promega). Lysates containing 30 μg of RNA were digested with P1 nuclease (30 U/μg RNA) for 1 h at 30°C with shaking at 250 rpm. Ribosome protected fragments were pelleted via sucrose cushion in an Optima TLX Ultracentrifuge with a TLA 110 rotor at 100,000 RPM for 1 h at 4°C. Pelleted RNA was size selected via mirRICH enrichment, as described^49,50^.

For cDNA synthesis and library preparation, 100–500 ng of mirRICH-purified RNA was used as the input for cDNA synthesis, using the improved ordered two-template relay protocol (OTTR v2^51^). Modifications to OTTR v1 include dilution of input RNA to 9 μL, followed by the sequential addition of 2 μL buffer 1A, 1 μL buffer 1B2, and 10μM TurboTail Pol (BoMoC W403AF753A). The reaction was incubated for 2 h at 30°C. Next, 1 μL of buffer 2 and 0.5 µL of rSAP (New England Biolabs, M0371S) premixed with 0.5 µL of 50% glycerol were added. After a 15 min incubation at 37°C, 1 µL of buffer 3 was added, followed by 5 min at 65°C, after which the reaction was placed on ice. Subsequently, 1 μL of buffer 4A, 1 µL of buffer 4B2, and 1 µL of 10 µM Relay Pol were added, followed by incubation for 30 min at 37°C. Reverse transcriptase activity was inactivated by a 5 min incubation at 70°C.

After cDNA synthesis, protein and RNA were removed by RNase A, RNase H, and proteinase K digestion and cDNA products were recovered by Zymo Oligo Clean & Concentrator (Zymo Research, D4061). cDNA was size selected on an 8% denaturing polyacrylamide gel, with imaging to detect IR800 as described^49^. cDNA was quantified using qPCR^48^ and libraries were constructed with 13 amplification cycles using Illumina dual indexing primers (UCSF CAT) and Q5 polymerase. PCR products were purified by Zymo DNA Clean and Concentrator columns followed by AMPure XP (Beckman, A63880) size selection. DNA libraries were pooled according to concentrations provided by TapeStation (Agilent). Libraries were sequenced on an Illumina NovaSeq X with single-end 100 bp reads (UCSF CAT). Sequencing data were processed using cutadapt to trim adapter sequences (constant AGATCGGAAGAGCACACGTCTGAACTCCAGTCAC and 7mer UMI) and deconvolve multiplexed libraries (doi://10.14806/ej.17.1.200). Trimmed reads were aligned by bowtie2 to remove rRNA reads followed by splice-aware genomic alignment of non-rRNA by hisat2. Translation efficiencies were determined with DESeq2.

#### Proteomics

4D-DIA proteomics analysis was performed by Creative Proteomics. Yeast cells were lysed in BPP buffer (100 mM EDTA, 50 mM borax, 50 mM vitamin C, 30% sucrose, 100 mM Tris-HCl, 1% Triton X-100, 1–5% PVPP, and 5 mM DTT, pH 8.0) supplemented with protease inhibitors. Cells were disrupted at −40 °C (4X120 s cycles), and proteins were extracted with Tris-saturated phenol (pH 8.0). After centrifugation, the phenolic phase was re-extracted with BPP buffer, and proteins were precipitated overnight with 0.1 M ammonium acetate in methanol at −20 °C. Pellets were washed with 90% acetone, air-dried, and dissolved in 8 M urea. Protein concentrations were determined using the BCA assay. Protein digestion was performed using the iST Sample Preparation Kit (PreOmics) following the manufacturer’s instructions. Briefly, samples were heated at 95 °C for 10 min, digested with trypsin at 37 °C for 2 h, and peptides were desalted and dried before LC–MS/MS analysis.

Peptides (200 ng) were analyzed on a timsTOF Pro2 mass spectrometer (Bruker Daltonik) coupled to a nanoElute 2 LC system using a C18 column (8 cm × 75 µm, 1.7 µm, IonOpticks). Separation was performed at 50 °C with a 60 min gradient (2.2–90% solvent B: 80% acetonitrile, 0.1% formic acid) at 300 nL/min. Data were acquired in diaPASEF mode (m/z 400–1,000, 25 × 25 Th windows, collision energy 59–20 eV). Raw DIA data were processed in Spectronaut v20.1 (Biognosys) against the *S. cerevisiae* UniProt database using default settings. Trypsin specificity, carbamidomethylation (C, fixed), oxidation (M), and N-terminal acetylation (variable) were applied. A 1% FDR was enforced at all levels, and quantification used the MaxLFQ algorithm. Proteins with fold-change > 1.5 (*p* < 0.05) were considered upregulated and < 1/1.5 (*p* < 0.05) downregulated. 3545 proteins were identified across all samples.

#### Cell culture and methionine starvation

HEK293T and Neuro-2a cells were maintained in a Heracell humidified incubator at 37°C with 5% CO₂. Both cell lines were cultured in high glucose DMEM (Sigma, D5796) supplemented with 10% fetal bovine serum (Sigma, F6178) and 1X Penicillin-Streptomycin (Gibco, 15140-122). Upon reaching ∼80% confluency, cells were washed three times with PBS and then incubated in DMEM (Gibco, 21013-024) containing 10% dialyzed FBS (Gibco, 26400-044), 1% Penicillin-Streptomycin (Gibco, 15140-122), GlutaMAX Supplement (Gibco, 35050-061; final concentration 2 mM), L-cysteine (Sigma, C7352; final concentration 0.2 mM), and with or without L-methionine (Sigma, M9625; final concentration 0.2 mM). After 18 h, cells were harvested, and total RNA was extracted using TRIzol reagent according to the manufacturer’s protocol.

#### *In vitro* transcription of capped RNAs

DNA template containing a T7 promoter was amplified by PCR. *In vitro* transcription was performed using T7 RNA polymerase (Promega, P2075) in a 50 µL reaction containing 1X Transcription Optimized Buffer (Promega, P2075), 10 mM DTT, 50 U rRNasin (Promega, N2511), 0.5 mM each of rATP, rUTP, and rCTP, 0.05 mM rGTP, 0.5 mM cap analog (m⁷GpppG or GpppG), 5 µg DNA template, and 40 U T7 RNA polymerase. The reaction was incubated at 37 °C for 1 h, after which an additional 40 U of T7 RNA polymerase was added and incubation continued for another 1 h. The reaction was then treated with DNase I and purified using the RNA Clean & Concentrator-25 kit (Zymo Research, R1017).

#### *In vitro* translation with yeast extract

Cell-free translation was performed using yeast extracts prepared as previously described^35^. Briefly, yeast cells were grown in YPD to mid-log phase (OD_600_ = 0.6-0.8) and harvested by centrifugation. Cell pellets were resuspended in buffer A (30 mM HEPES-KOH pH 7.4, 100 mM KOAc, 3 mM Mg(OAc)_2_, and 1 mM DTT) with 85 g/L mannitol at 2:3 volume (ml):pellet weight (g) ratio and lysed using a cryomill. Lysates were centrifuged at 30,000 x g for 15 min at 4 °C. The middle fraction was transferred into a new tube and centrifuged again at 100,000 x g for 35 min at 4°C. The middle fraction was collected and desalted into buffer A with 20% glycerol using PD-10 columns (Cytiva, 17085101) while monitoring RNA concentrations. RNA-rich fractions were collected and flash-frozen in liquid nitrogen.

*In vitro* translation reactions were assembled in 15 µL volume with 7.5 µL of yeast translation extract, 250 ng of *in vitro* transcribed RNA, 50 mM KOAc, 2 mM Mg(OAc)_2_, 10 μM of amino acid mix, 0.4 U/μL RiboLock (ThermoFisher, EO0381), 0.06 U/μL creatine kinase (BioVision, P1301), and 1X energy mix containing 20 mM HEPES (pH 7.6), 1 mM ATP, 0.1 mM GTP, 20 mM creatine phosphate, and 2 mM DTT. Reactions were incubated at room temperature for 2 h and stopped by adding SDS loading dye. Translation efficiency was assessed by Western blot.

#### *In vivo* translation in yeast spheroplasts

Yeast spheroplasts were prepared for electroporation as previously described^52^ with modifications. Cells grown in YPL medium were diluted to an OD_600_ of 0.008 in 50 mL YPL and cultured overnight. Cultures were harvested at OD_600_ = 0.7–0.8 and resuspended in spheroplast buffer (50 mM Tris-HCl, pH 7.4, 1 mM MgCl_2_, 30 mM DTT, 15 mM β-mercaptoethanol, 1 M sorbitol). Spheroplasts were generated by treatment with 200 µL of 1 mg/mL Zymolyase 100T for 15–20 min at 30°C. Unless otherwise indicated, spheroplasts were centrifuged at 1,000 × *g* for 5 min for all subsequent washes. Following two washes with spheroplast buffer, spheroplasts were resuspended in 1 mL spheroplast buffer, diluted into 9 mL YPL supplemented with 1 M sorbitol, and allowed to recover for 90 min at 30°C with gentle shaking. Cells were then washed twice with 1 M sorbitol and resuspended in ice-cold 1 M sorbitol.

For electroporation, 180 µL of spheroplasts were mixed with 2 µg of *in vitro*-transcribed RNA in pre-chilled 0.2-cm gap cuvettes (Bio-Rad, 1652086) and electroporated using a Gene Pulser Xcell (Bio-Rad) at 800 V, 25 µF, and 1,000 Ω (time constant, 23–25 ms). Immediately after electroporation, 1.2 mL of ice-cold YPL or SL medium supplemented with 1 M sorbitol was added, and cells were recovered for 2 h at 30°C with gentle shaking. Cells were then harvested, flash-frozen in liquid nitrogen, and stored at −80°C until analysis.

NanoLuc luciferase activity was measured using the Nano-Glo Luciferase Assay System (Promega, N1110). Frozen spheroplast pellets were lysed in 100 µL of 1× Passive Lysis Buffer (Promega, E1941) by vortexing at 2,000 rpm for 15 min. Lysates were clarified by centrifugation at maximum speed for 5 min, and 50 µL of the supernatant was mixed with 100 µL of Nano-Glo reagent in a white 96-well plate. Following incubation at room temperature, luminescence was measured using a Cytation 5 plate reader (BioTek). Luciferase activity was normalized to total protein concentration determined by the Bradford assay (Bio-Rad, 5000001).

#### Biotinylated RNA pull-down

0 mL of yeast cells were grown to OD_600_ ∼ 0.8 and collected by centrifugation. Cells were lysed by bead-beating 3X 30 s in lysis buffer containing 20 mM Tris-HCl pH=7.5, 100 mM KCl, 5 mM MgCl_2_, and 0.5% Nodidet P-40, 1 mM NaVO3, 5 mM NaF, and 200 U/ml rRNasin. Lysate was centrifuged at 8000 xg for 5 min at 4 °C. Supernatant was collected and normalized by total protein concentration. 50 µL of Streptavidin beads was washed in 1X wash buffer (10 mM Tris-HCl pH=8.0, 250 mM NaCl, 0.05% (v/v) Triton X-100, 2 mM MgCl2) twice and incubated with 1 µg of biotinylated RNA oligo at RT for 10 min and washed with wash buffer twice. The pull-down reaction consists of RNA-bound beads from above, 500 µg of cell lysate in 250 µL lysis buffer, 1X protease inhibitor cocktail (Roche), 100 U of rRNasin and 250 µL of 2X wash buffer (20 mM Tris-HCl pH=8.0, 500 mM NaCl, 0.1% (v/v) Triton X-100, 4 mM MgCl_2_). The reaction was incubated by rotating in room temperature for 40 min and washed 3X with wash buffer. Washed beads were resuspended in a denaturing loading dye for NuPAGE gels and heated at 95°C for 5 min before being subjected to western blot analysis. For proteomics, beads were further washed in 1X PBS for three times and submitted to UC Davis Proteomics core for digestion and data-independent acquisition (DIA) Profiling.

## Statistical analysis

Statistical analyses of bar graphs were conducted in Excel using the two-tailed, unpaired Student’s t-test assuming equal variance. *, **, *** indicate p < 0.05, p < 0.01, < 0.001, respectively. Pathway enrichment analysis was conducted using YeastEnrichr^53^ and Enrichr^53^. Overrepresentation of gene sets was assessed by Fisher’s exact test, and p values were adjusted for multiple hypothesis testing using the Benjamini–Hochberg false discovery rate procedure.

## Declaration of generative AI and AI-assisted technologies in the writing process

Claude was used during the preparation of this manuscript to assist in generating the original figures for Fig. 6C and 6F. These were reviewed and revised by the authors, who take full responsibility for the final content of the published article.

## References

1. Zheng, D., Wang, R., Ding, Q., Wang, T., Xie, B., Wei, L., Zhong, Z., and Tian, B. (2018). Cellular stress alters 3’UTR landscape through alternative polyadenylation and isoform-specific degradation. Nat Commun 9, 2268. 10.1038/s41467-018-04730-7.

2. Woo, Y.M., Kwak, Y., Namkoong, S., Kristjánsdóttir, K., Lee, S.H., Lee, J.H., and Kwak, H. (2018). TED-Seq Identifies the Dynamics of Poly(A) Length during ER Stress. Cell Rep 24, 3630–3641.e3637. 10.1016/j.celrep.2018.08.084.

3. Liu, Y., Zhao, H., Shao, F., Zhang, Y., Nie, H., Zhang, J., Li, C., Hou, Z., Chen, Z.J., Wang, J., et al. (2023). Remodeling of maternal mRNA through poly(A) tail orchestrates human oocyte-to-embryo transition. Nat Struct Mol Biol 30, 200–215. 10.1038/s41594-022-00908-2.

4. Pendleton, K.E., Chen, B., Liu, K., Hunter, O.V., Xie, Y., Tu, B.P., and Conrad, N.K. (2017). The U6 snRNA m(6)A Methyltransferase METTL16 Regulates SAM Synthetase Intron Retention. Cell 169, 824–835.e814. 10.1016/j.cell.2017.05.003.

5. Mattay, J. (2022). Noncanonical metabolite RNA caps: Classification, quantification, (de)capping, and function. Wiley Interdiscip Rev RNA 13, e1730. 10.1002/wrna.1730.

6. Wang, J., Alvin Chew, B.L., Lai, Y., Dong, H., Xu, L., Balamkundu, S., Cai, W.M., Cui, L., Liu, C.F., Fu, X.Y., et al. (2019). Quantifying the RNA cap epitranscriptome reveals novel caps in cellular and viral RNA. Nucleic Acids Res 47, e130. 10.1093/nar/gkz751.

7. Marcotrigiano, J., Gingras, A.C., Sonenberg, N., and Burley, S.K. (1997). Cocrystal structure of the messenger RNA 5’ cap-binding protein (eIF4E) bound to 7-methyl-GDP. Cell 89, 951–961. 10.1016/s0092-8674(00)80280-9.

8. Xing, Z., and Tu, B.P. (2025). Mechanisms and rationales of SAM homeostasis. Trends Biochem Sci 50, 242–254. 10.1016/j.tibs.2024.12.009.

9. Schwer, B., Saha, N., Mao, X., Chen, H.W., and Shuman, S. (2000). Structure-function analysis of yeast mRNA cap methyltransferase and high-copy suppression of conditional mutants by AdoMet synthase and the ubiquitin conjugating enzyme Cdc34p. Genetics 155, 1561–1576.

10. Furuichi, Y., Morgan, M., Shatkin, A.J., Jelinek, W., Salditt-Georgieff, M., and Darnell, J.E. (1975). Methylated, blocked 5 termini in HeLa cell mRNA. Proc Natl Acad Sci U S A 72, 1904–1908. 10.1073/pnas.72.5.1904.

11. Cole, M.D., and Cowling, V.H. (2009). Specific regulation of mRNA cap methylation by the c-Myc and E2F1 transcription factors. Oncogene 28, 1169–1175. 10.1038/onc.2008.463.

12. Aregger, M., and Cowling, V.H. (2012). E2F1-dependent methyl cap formation requires RNA pol II phosphorylation. Cell Cycle 11, 2146–2148. 10.4161/cc.20620.

13. Fernandez-Sanchez, M.E., Gonatopoulos-Pournatzis, T., Preston, G., Lawlor, M.A., and Cowling, V.H. (2009). S-adenosyl homocysteine hydrolase is required for Myc-induced mRNA cap methylation, protein synthesis, and cell proliferation. Mol Cell Biol 29, 6182–6191. 10.1128/mcb.00973-09.

14. Aregger, M., Kaskar, A., Varshney, D., Fernandez-Sanchez, M.E., Inesta-Vaquera, F.A., Weidlich, S., and Cowling, V.H. (2016). CDK1-Cyclin B1 Activates RNMT, Coordinating mRNA Cap Methylation with G1 Phase Transcription. Mol Cell 61, 734–746. 10.1016/j.molcel.2016.02.008.

15. Zhang, L.S., Liu, C., Ma, H., Dai, Q., Sun, H.L., Luo, G., Zhang, Z., Zhang, L., Hu, L., Dong, X., and He, C. (2019). Transcriptome-wide Mapping of Internal N(7)-Methylguanosine Methylome in Mammalian mRNA. Mol Cell 74, 1304–1316.e1308. 10.1016/j.molcel.2019.03.036.

16. Galloway, A., Kaskar, A., Ditsova, D., Atrih, A., Yoshikawa, H., Gomez-Moreira, C., Suska, O., Warminski, M., Grzela, R., Lamond, A.I., et al. (2021). Upregulation of RNA cap methyltransferase RNMT drives ribosome biogenesis during T cell activation. Nucleic Acids Res 49, 6722–6738. 10.1093/nar/gkab465.

17. Knop, K., Gomez-Moreira, C., Galloway, A., Ditsova, D., and Cowling, V.H. (2024). RAM is upregulated during T cell activation and is required for RNA cap formation and gene expression. Discov Immunol 3, kyad021. 10.1093/discim/kyad021.

18. Jiao, X., Xiang, S., Oh, C., Martin, C.E., Tong, L., and Kiledjian, M. (2010). Identification of a quality-control mechanism for mRNA 5’-end capping. Nature 467, 608–611. 10.1038/nature09338.

19. Chang, J.H., Jiao, X., Chiba, K., Oh, C., Martin, C.E., Kiledjian, M., and Tong, L. (2012). Dxo1 is a new type of eukaryotic enzyme with both decapping and 5’-3’ exoribonuclease activity. Nat Struct Mol Biol 19, 1011–1017. 10.1038/nsmb.2381.

20. Jiao, X., Chang, J.H., Kilic, T., Tong, L., and Kiledjian, M. (2013). A mammalian pre-mRNA 5’ end capping quality control mechanism and an unexpected link of capping to pre-mRNA processing. Mol Cell 50, 104–115. 10.1016/j.molcel.2013.02.017.

21. Sutter, B.M., Wu, X., Laxman, S., and Tu, B.P. (2013). Methionine inhibits autophagy and promotes growth by inducing the SAM-responsive methylation of PP2A. Cell 154, 403–415. 10.1016/j.cell.2013.06.041.

22. Malabat, C., Feuerbach, F., Ma, L., Saveanu, C., and Jacquier, A. (2015). Quality control of transcription start site selection by nonsense-mediated-mRNA decay. Elife 4. 10.7554/eLife.06722.

23. Mao, X., Schwer, B., and Shuman, S. (1995). Yeast mRNA cap methyltransferase is a 50-kilodalton protein encoded by an essential gene. Mol Cell Biol 15, 4167–4174. 10.1128/mcb.15.8.4167.

24. Morawska, M., and Ulrich, H.D. (2013). An expanded tool kit for the auxin-inducible degron system in budding yeast. Yeast 30, 341–351. 10.1002/yea.2967.

25. Ye, C., Sutter, B.M., Wang, Y., Kuang, Z., and Tu, B.P. (2017). A Metabolic Function for Phospholipid and Histone Methylation. Mol Cell 66, 180–193.e188. 10.1016/j.molcel.2017.02.026.

26. Ye, C., Sutter, B.M., Wang, Y., Kuang, Z., Zhao, X., Yu, Y., and Tu, B.P. (2019). Demethylation of the Protein Phosphatase PP2A Promotes Demethylation of Histones to Enable Their Function as a Methyl Group Sink. Mol Cell 73, 1115–1126.e1116. 10.1016/j.molcel.2019.01.012.

27. Trcek, T., Larson, D.R., Moldón, A., Query, C.C., and Singer, R.H. (2011). Single-molecule mRNA decay measurements reveal promoter-regulated mRNA stability in yeast. Cell 147, 1484–1497. 10.1016/j.cell.2011.11.051.

28. Willnow, S., Martin, M., Lüscher, B., and Weinhold, E. (2012). A selenium-based click AdoMet analogue for versatile substrate labeling with wild-type protein methyltransferases. Chembiochem 13, 1167–1173. 10.1002/cbic.201100781.

29. Gopalakrishnan, R., and Winston, F. (2021). The histone chaperone Spt6 is required for normal recruitment of the capping enzyme Abd1 to transcribed regions. J Biol Chem 297, 101205. 10.1016/j.jbc.2021.101205.

30. Ye, C., and Tu, B.P. (2018). Sink into the Epigenome: Histones as Repositories That Influence Cellular Metabolism. Trends Endocrinol Metab 29, 626–637. 10.1016/j.tem.2018.06.002.

31. Strahl, B.D., Grant, P.A., Briggs, S.D., Sun, Z.W., Bone, J.R., Caldwell, J.A., Mollah, S., Cook, R.G., Shabanowitz, J., Hunt, D.F., and Allis, C.D. (2002). Set2 is a nucleosomal histone H3-selective methyltransferase that mediates transcriptional repression. Mol Cell Biol 22, 1298–1306. 10.1128/mcb.22.5.1298-1306.2002.

32. Maeda, T., Takekawa, M., and Saito, H. (1995). Activation of yeast PBS2 MAPKK by MAPKKKs or by binding of an SH3-containing osmosensor. Science 269, 554–558. 10.1126/science.7624781.

33. Tutucci, E., Vera, M., Biswas, J., Garcia, J., Parker, R., and Singer, R.H. (2018). An improved MS2 system for accurate reporting of the mRNA life cycle. Nat Methods 15, 81–89. 10.1038/nmeth.4502.

34. Chassé, H., Boulben, S., Costache, V., Cormier, P., and Morales, J. (2017). Analysis of translation using polysome profiling. Nucleic Acids Res 45, e15. 10.1093/nar/gkw907.

35. Trainor, B.M., Komar, A.A., Pestov, D.G., and Shcherbik, N. (2021). Cell-free Translation: Preparation and Validation of Translation-competent Extracts from Saccharomyces cerevisiae. Bio Protoc 11, e4093. 10.21769/BioProtoc.4093.

36. Gillian-Daniel, D.L., Gray, N.K., Aström, J., Barkoff, A., and Wickens, M. (1998). Modifications of the 5’ cap of mRNAs during Xenopus oocyte maturation: independence from changes in poly(A) length and impact on translation. Mol Cell Biol 18, 6152–6163. 10.1128/mcb.18.10.6152.

37. Chen, Y., Cai, H., Pan, J., Xiang, N., Tien, P., Ahola, T., and Guo, D. (2009). Functional screen reveals SARS coronavirus nonstructural protein nsp14 as a novel cap N7 methyltransferase. Proc Natl Acad Sci U S A 106, 3484–3489. 10.1073/pnas.0808790106.

38. Galloway, A., and Cowling, V.H. (2019). mRNA cap regulation in mammalian cell function and fate. Biochim Biophys Acta Gene Regul Mech 1862, 270–279. 10.1016/j.bbagrm.2018.09.011.

39. Liu, K., Santos, D.A., Hussmann, J.A., Wang, Y., Sutter, B.M., Weissman, J.S., and Tu, B.P. (2021). Regulation of translation by methylation multiplicity of 18S rRNA. Cell Rep 34, 108825. 10.1016/j.celrep.2021.108825.

40. Enroth, C., Poulsen, L.D., Iversen, S., Kirpekar, F., Albrechtsen, A., and Vinther, J. (2019). Detection of internal N7-methylguanosine (m7G) RNA modifications by mutational profiling sequencing. Nucleic Acids Res 47, e126. 10.1093/nar/gkz736.

41. Muthukrishnan, S., Both, G.W., Furuichi, Y., and Shatkin, A.J. (1975). 5′-Terminal 7-methylguanosine in eukaryotic mRNA is required for translation. Nature 255, 33–37. 10.1038/255033a0.

42. Trotman, J.B., and Schoenberg, D.R. (2019). A recap of RNA recapping. Wiley Interdiscip Rev RNA 10, e1504. 10.1002/wrna.1504.

43. van Dijken, J.P., Bauer, J., Brambilla, L., Duboc, P., Francois, J.M., Gancedo, C., Giuseppin, M.L., Heijnen, J.J., Hoare, M., Lange, H.C., et al. (2000). An interlaboratory comparison of physiological and genetic properties of four Saccharomyces cerevisiae strains. Enzyme Microb Technol 26, 706–714. 10.1016/s0141-0229(00)00162-9.

44. Longtine, M.S., McKenzie, A., 3rd, Demarini, D.J., Shah, N.G., Wach, A., Brachat, A., Philippsen, P., and Pringle, J.R. (1998). Additional modules for versatile and economical PCR-based gene deletion and modification in Saccharomyces cerevisiae. Yeast 14, 953–961. 10.1002/(sici)1097-0061(199807)14:10<953::Aid-yea293>3.0.Co;2-u.

45. Veatch, J.R., McMurray, M.A., Nelson, Z.W., and Gottschling, D.E. (2009). Mitochondrial dysfunction leads to nuclear genome instability via an iron-sulfur cluster defect. Cell 137, 1247–1258. 10.1016/j.cell.2009.04.014.

46. Liu, K., Sutter, B.M., and Tu, B.P. (2021). Autophagy sustains glutamate and aspartate synthesis in Saccharomyces cerevisiae during nitrogen starvation. Nat Commun 12, 57. 10.1038/s41467-020-20253-6.

47. Josefsen, K., and Nielsen, H. (2011). Northern blotting analysis. Methods Mol Biol 703, 87–105. 10.1007/978-1-59745-248-9_7.

48. McGlincy, N.J., and Ingolia, N.T. (2017). Transcriptome-wide measurement of translation by ribosome profiling. Methods 126, 112–129. 10.1016/j.ymeth.2017.05.028.

49. Ferguson, L., Upton, H.E., Pimentel, S.C., Mok, A., Lareau, L.F., Collins, K., and Ingolia, N.T. (2023). Streamlined and sensitive mono- and di-ribosome profiling in yeast and human cells. Nat Methods 20, 1704–1715. 10.1038/s41592-023-02028-1.

50. Choi, C., Yoon, S., Moon, H., Bae, Y.U., Kim, C.B., Diskul-Na-Ayudthaya, P., Ngu, T.V., Munir, J., Han, J., Park, S.B., et al. (2018). mirRICH, a simple method to enrich the small RNA fraction from over-dried RNA pellets. RNA Biol 15, 763–772. 10.1080/15476286.2018.1451723.

51. Ferguson, L., Upton, H.E., Pimentel, S.C., Jeans, C., Ingolia, N.T., and Collins, K. (2025). Improved precision, sensitivity, and adaptability of ordered two-template relay cDNA library preparation for RNA sequencing. Rna 31, 224–244. 10.1261/rna.080318.124.

52. Searfoss, A.M., Masison, D.C., and Wickner, R.B. (2002). Protein synthesis assayed by electroporation of mRNA in Saccharomyces cerevisiae. Methods Enzymol 351, 631–639. 10.1016/s0076-6879(02)51873-5.

53. Kuleshov, M.V., Jones, M.R., Rouillard, A.D., Fernandez, N.F., Duan, Q., Wang, Z., Koplev, S., Jenkins, S.L., Jagodnik, K.M., Lachmann, A., et al. (2016). Enrichr: a comprehensive gene set enrichment analysis web server 2016 update. Nucleic Acids Res 44, W90–97. 10.1093/nar/gkw377.

54. Liu, Y., and Xie, J. (2020). Cauchy combination test: a powerful test with analytic p-value calculation under arbitrary dependency structures. J Am Stat Assoc 115, 393–402. 10.1080/01621459.2018.1554485.

