## Supplemental Figures 1-6 for "Metabolic stress reveals widespread accumulation of cap-unmethylated RNAs"

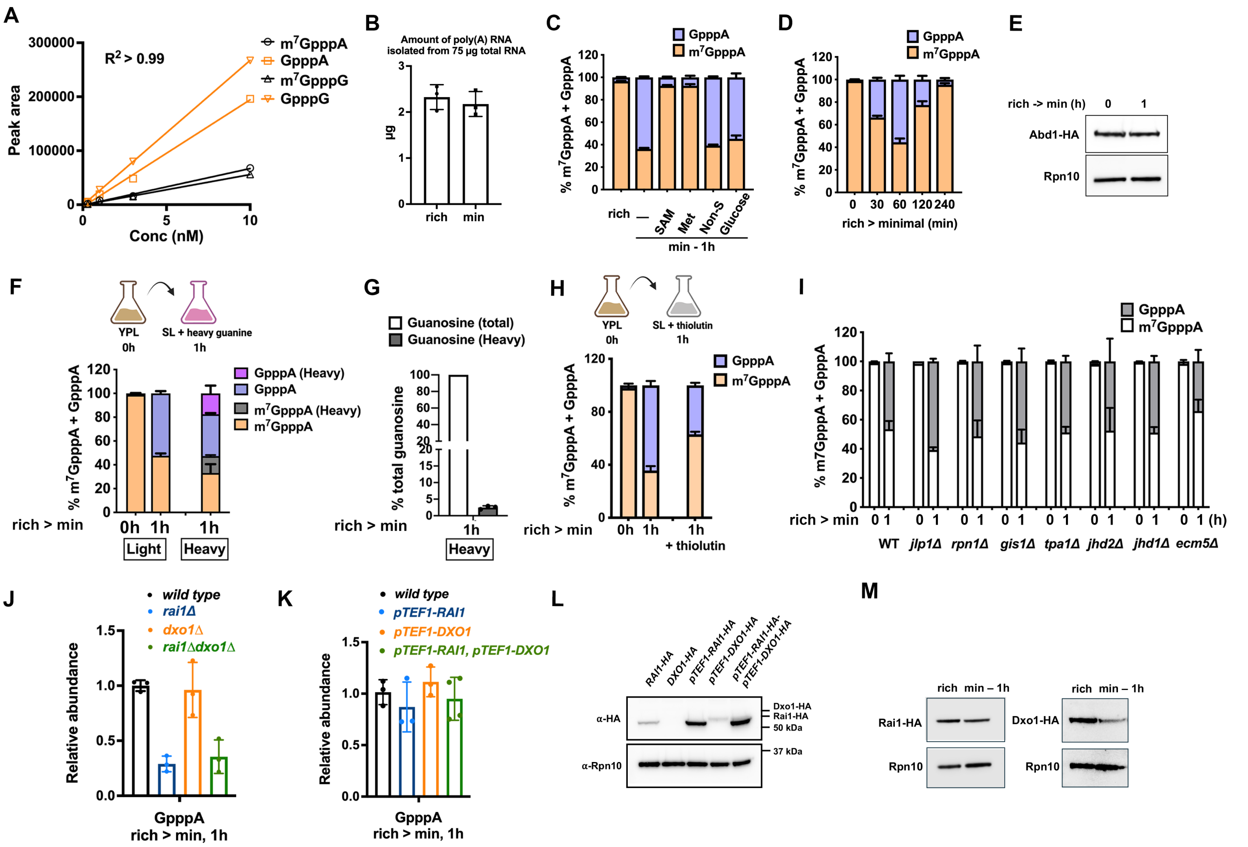


**Fig. S1. (A)** Representative standard curves generated using purified cap analogs. **(B)** Quantity of poly(A)+ mRNA isolated from 75 µg of total RNA extracted from cells grown in rich or minimal medium. **(C)** LC-MS/MS analysis of cap dinucleotide percentages with the addition of 0.5 mM SAM, methionine, non-sulfur amino acid mix, or 2% glucose during a 1 h switch from rich to minimal medium. **(D)** LC-MS/MS analysis of cap dinucleotide percentages after switching cells from rich to minimal medium for indicated time. **(E)** Western blot analysis of Abd1 levels in yeast cells following a switch from rich medium to minimal medium. **(F)** LC-MS/MS analysis of cap dinucleotide percentages with the addition of heavy guanine during a 1 h switch from rich to minimal medium. **(G)** Percentage of heavy-labeled guanosine as compared to light guanosine when cells were grown in minimal medium containing heavy guanine for 1 h. **(H)** LC-MS/MS analysis of cap dinucleotide percentage with the addition of thiolutin (20 µg/ml) when switching cells from rich to minimal medium for 1 h. **(I)** LC-MS/MS analysis of cap dinucleotide percentages in wild type and demethylase deletion strains. **(J, K)** Relative abundance of GpppA cap structures of cells grown in minimal medium for 1 h. Data were measured by LC-MS/MS. **(L)** Western blot analysis of strains from (K). **(M)** Western blot analysis of Rai1 and Dxo1 protein levels following a shift to minimal medium. All bar graph data represent ≥ 3 biological replicates and are shown as mean ± SD.

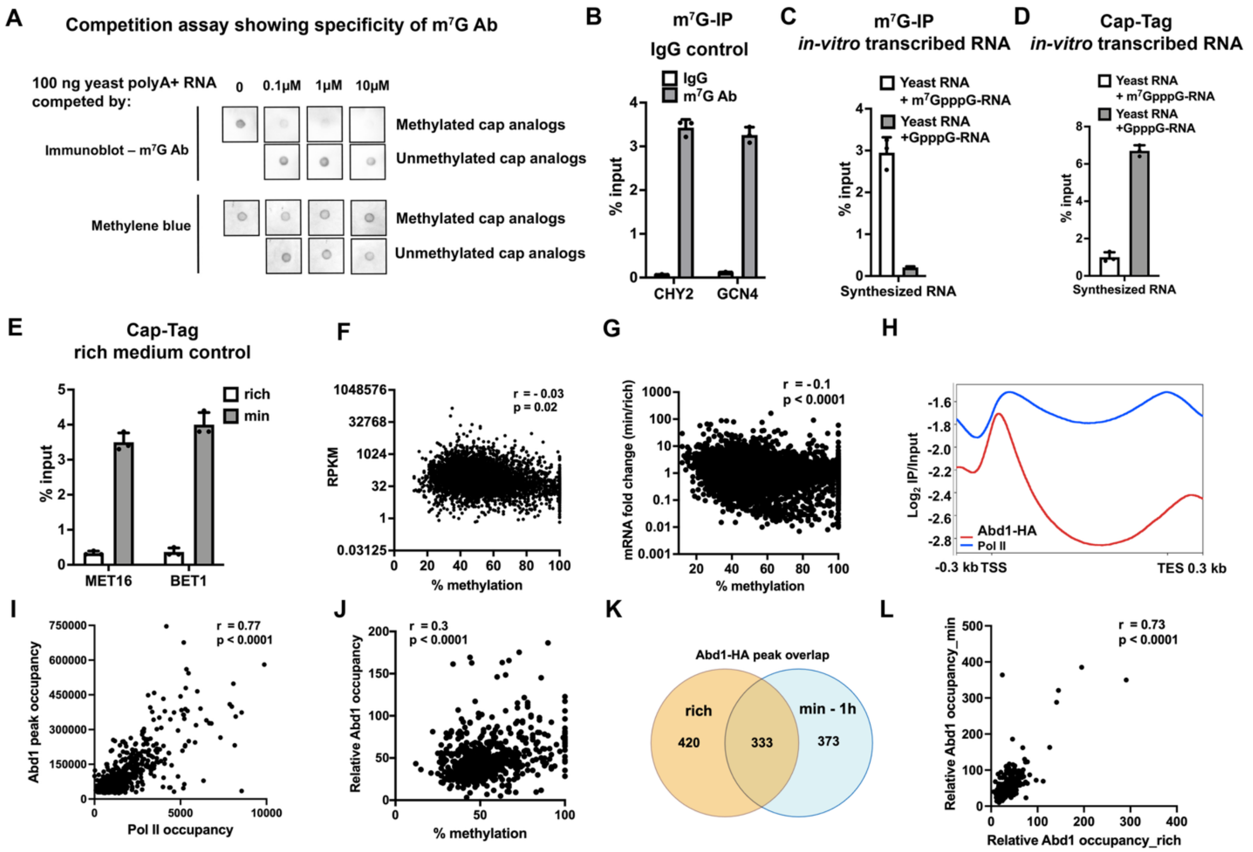


**Fig. S2. (A)** Dot blot analysis of $m^7G$ cap in yeast mRNAs extracted from cells grown in rich medium, using either a mixture of methylated cap analogs ($m^7$GpppA and $m^7$GpppG) or unmethylated cap analogs (GpppA and GpppG) as competitors for antibody binding. **(B)** Comparison of $m^7G$-IP-qPCR results for mRNAs extracted from cells grown in rich medium, using IgG as a control. Percent input was measured for two randomly selected mRNAs, CHY2 and GCN4. **(C)** $m^7G$-IP-qPCR results of *in vitro* transcribed and capped RNAs mixed with 100 ng of yeast mRNAs. **(D)** Cap-Tag-qPCR results of *in vitro* transcribed and capped RNAs mixed with 100 ng of yeast mRNAs. **(E)** Cap-Tag-qPCR results of mRNAs extracted from cells grown in rich or minimal media. Percent input was measured for two lowly-methylated mRNAs identified in the $m^7G$-IP experiment. **(F, G)** Correlation of $m^7G$ methylation levels (measured by $m^7G$-IP) with (F) Reads Per Kilobase of transcript per Million mapped reads (RPKM), (G) fold change in mRNA levels between minimal and rich media. **(H)** Metagene profiles showing average Abd1-HA and Rpb1 (Pol II) occupancy in cells grown in minimal medium for 1 h. **(I)** Correlation between Abd1 and Rpb1 (Pol II) occupancy. **(J)** Correlation between relative Abd1 occupancy (normalized to RNA polymerase II occupancy) and mRNA cap methylation. **(K)** Overlap of Abd1-HA ChIP-seq peaks identified in cells grown in rich or minimal medium. **(L)** Correlation of relative Abd1 occupancy in cells grown in rich versus minimal medium. All bar graph data represent ≥ 3 biological replicates and are shown as mean ± SD.

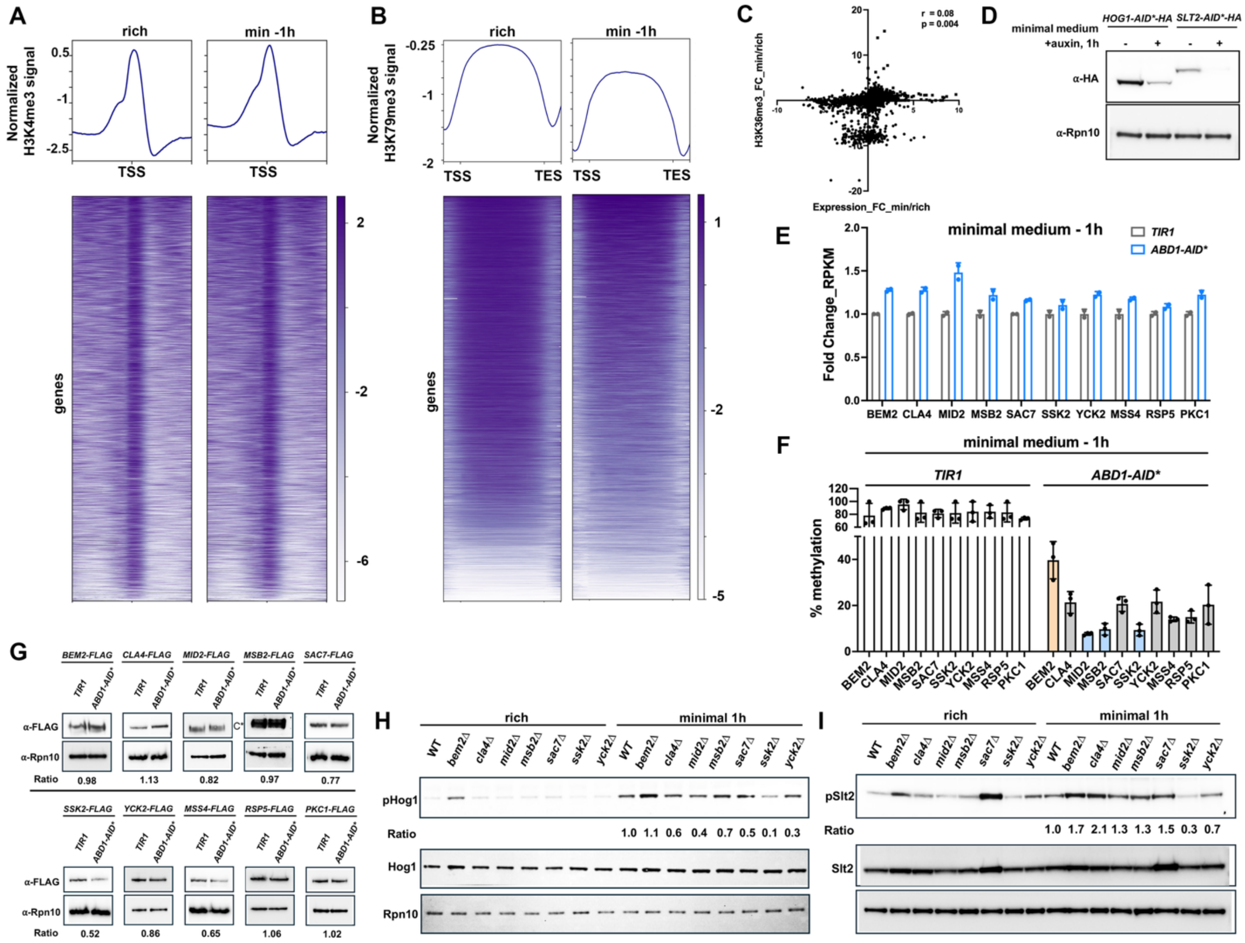

**Fig. S3. (A, B)** ChIP-seq profiles displaying H3K4me3 (A) and H3K79me3 (B) distribution in cells grown in rich or minimal medium for 1 h, normalized to total H3 ChIP signals. **(C)** Correlation between changes in H3K36me3 levels and changes in mRNA expression levels (RPKM) after switching cells from rich to minimal medium for 1 h. **(D)** Western blot showing depletion of Hog1 and Slt2 using the AID* system. **(E)** Fold change in RPKM of MAPK pathway genes in *TIR1 and ABD1-AID** strains grown in minimal medium containing 200 µM auxin for 1 h; mean ± SD, *n* = 2. **(F)** Percent cap methylation of MAPK pathway genes in *TIR1* and *ABD1-AID** strains grown in minimal medium containing 200 µM auxin for 1 h, measured by m$^7$G-IP; mean ± SD, *n* = 3. **(G)** Western blot analysis of MAPK pathway proteins in *TIR1* and *ABD1-AID** strains grown in minimal medium containing 200 µM auxin for 1 h. Ratios indicate the abundance of each target protein in the *ABD1-AID** strain relative to the *TIR1* strain after normalization to Rpn10. *C indicates that only the cleaved form of Msb2 was detected. **(H-I)** Western blot analysis of phosphorylated and total Hog1 (H) and Slt2 (I) in single deletion strains of non-essential MAPK pathway genes. Rpn10 was used as a loading control. The ratio represents Rpn10-normalized p-Hog1 or p-Slt2 abundance in each deletion strain.

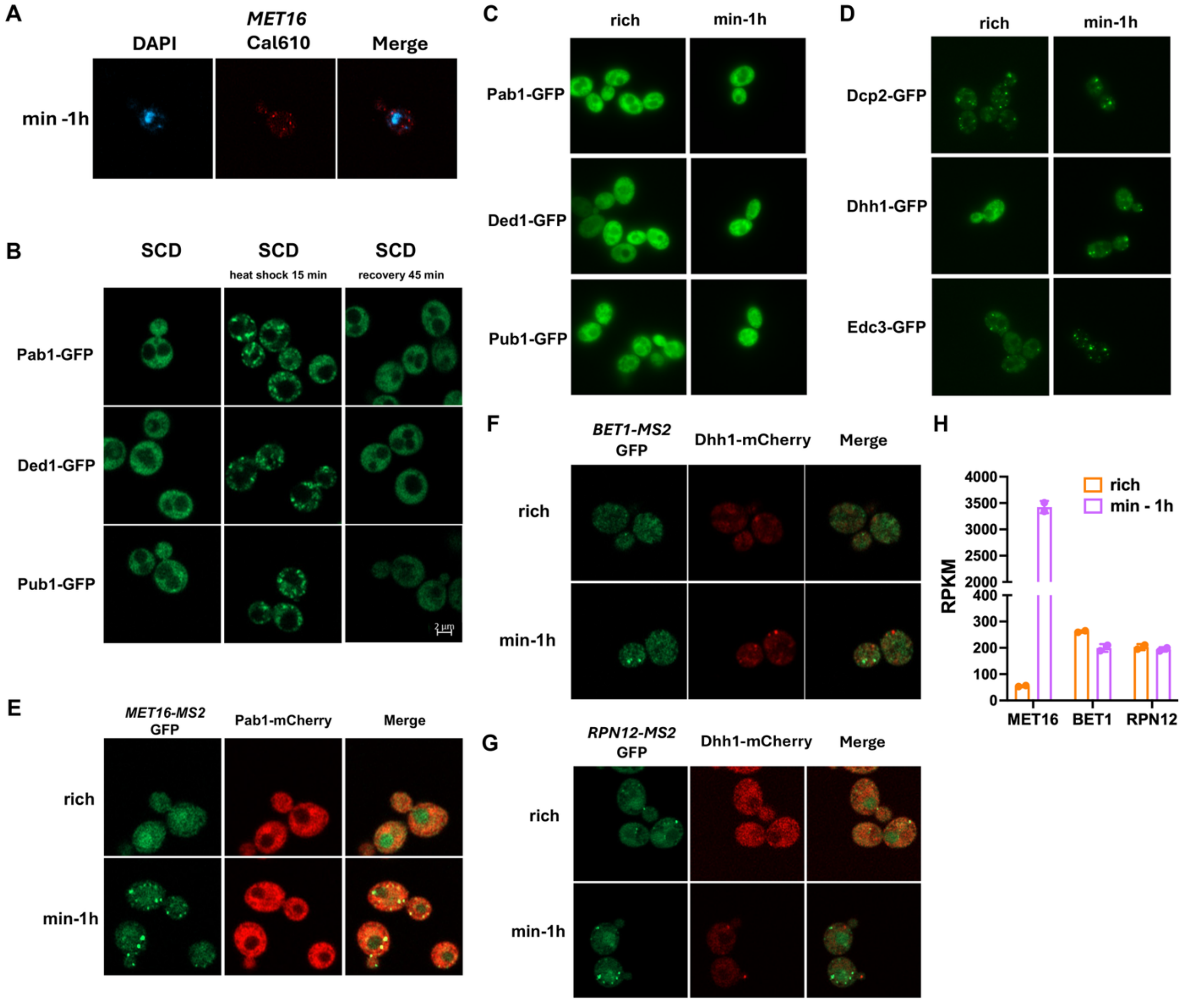


**Fig. S4.** **(A)** Microscopy images of yeast cells after a 1 h shift to minimal medium. Blue indicates DAPI staining of DNA; red indicates RNA FISH signal for *MET16* detected with Cal Fluor Red 610 dye. **(B)** Microscopy images of yeast strains expressing GFP-tagged stress granule markers including Pab1, Ded1, and Pub1, following heat shock at 42°C for 15 min and recovery for 45 min in synthetic complete medium containing 2% glucose; scale bar, 2 µm. **(C)** Microscopy images of GFP-tagged yeast stress granule markers before and after a 1 h shift to minimal medium. **(D)** Microscopy images of GFP-tagged yeast P-body markers including Dcp2, Dhh1, and Edc3, before and after a 1 h switch to minimal medium. **(E)** Microscopy images of MET16 mRNA (green) and stress granule marker Pab1-mCherry (red) before and after a 1 h shift to minimal medium. **(F, G)** Microscopy images of BET1 mRNA (F, % methylation = 33, green) or RPN12 mRNA (G, % methylation = 29, green); red indicates the P-body marker Dhh1-mCherry. **(H)** Transcript levels of *MET16*, *BET1*, and *RPN12*; mean ± SD, *n* = 2.

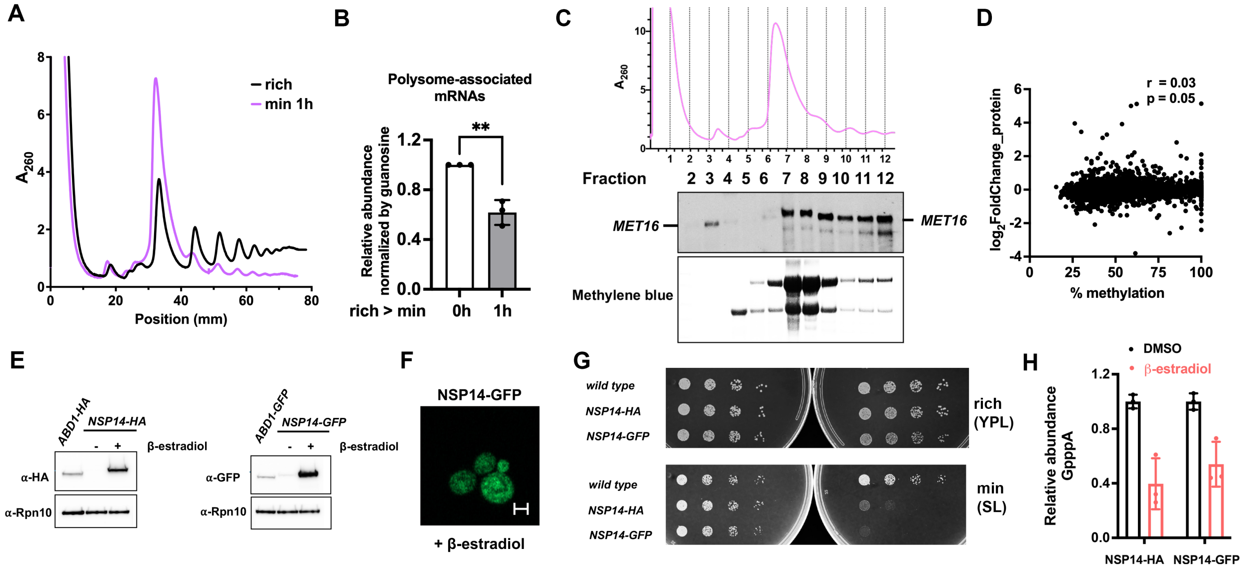


I

| KEGG pathway | $P_{adj}$ | Size |
|---|---|---|
| Cell cycle | 4 e-15 | 27/126 |
| Protein processing in ER | 6 e-13 | 21/88 |
| Meiosis | 9 e-09 | 20/130 |
| Glycolysis / Gluconeogenesis | 2 e-08 | 13/54 |
| RNA degradation | 2 e-08 | 14/62 |
| MAPK signaling pathway | 2 e-08 | 18/114 |
| Fatty acid degradation | 2 e-07 | 8/19 |

J

| KEGG pathway | $P_{adj}$ | Size |
|---|---|---|
| Spliceosome | 2 e-09 | 9/73 |
| Proteasome | 4 e-09 | 6/35 |
| RNA polymerase | 5 e-08 | 6/30 |
| RNA degradation | 0.001 | 4/62 |
| Ribosome biogenesis in eukaryotes | 0.002 | 4/75 |

**Fig. S5. (A)** Polysome profiling of cells grown in rich or minimal medium for 1 h, showing reduced polysome/monosome ratio in minimal medium (pink). **(B)** LC–MS/MS quantification of cap dinucleotides from mRNAs with adenosine or guanosine as the first transcribed nucleotide in pooled polysome fractions. **(C)** Northern blot analysis of *MET16* mRNA (upper band) using RNA extracted from polysome fractions. Polysome profiling was performed on cells grown in minimal medium for 1 h. Equal proportions of each fraction were loaded, and RNA concentrations were not normalized. MET16 mRNA is enriched in translating fractions: ~85% in monosome plus polysome fractions and ~70% in polysomes alone. **(D)** Correlation of $\log_2$FC of protein levels (min 1h/rich) and % methylation levels of each mRNA in cells grown in minimal medium for 1 h. **(E)** Western blot showing overexpression of Nsp14-HA and Nsp14-GFP following induction with 1 µM β-estradiol for 1 h in minimal medium. Rpn10 was used as a loading control. **(F)** Microscopy image of yeast cells expressing NSP14-GFP. Scale bar indicates 2 µm. **(G)** Serial dilution assays of wild type, *NSP14-HA*, and *NSP14-GFP* strains spotted on rich or minimal media supplemented with DMSO or 1 µM β-estradiol. **(H)** LC-MS/MS analysis of relative GpppA mRNA levels in *NSP14-HA* and *NSP14-GFP* cells with or without 1 µM β-estradiol. **(I)** KEGG pathway enrichment analysis of the yeast genes with conserved human orthologs and high cap methylation, corresponding to the overlap shown in Fig. 6D. **(J)** KEGG pathway enrichment analysis of the yeast genes with conserved human orthologs and low cap methylation, corresponding to the overlap shown in Fig. 6E. All bar graph data represent ≥ 3 biological replicates and are shown as mean ± SD. ** indicates $p < 0.01$.

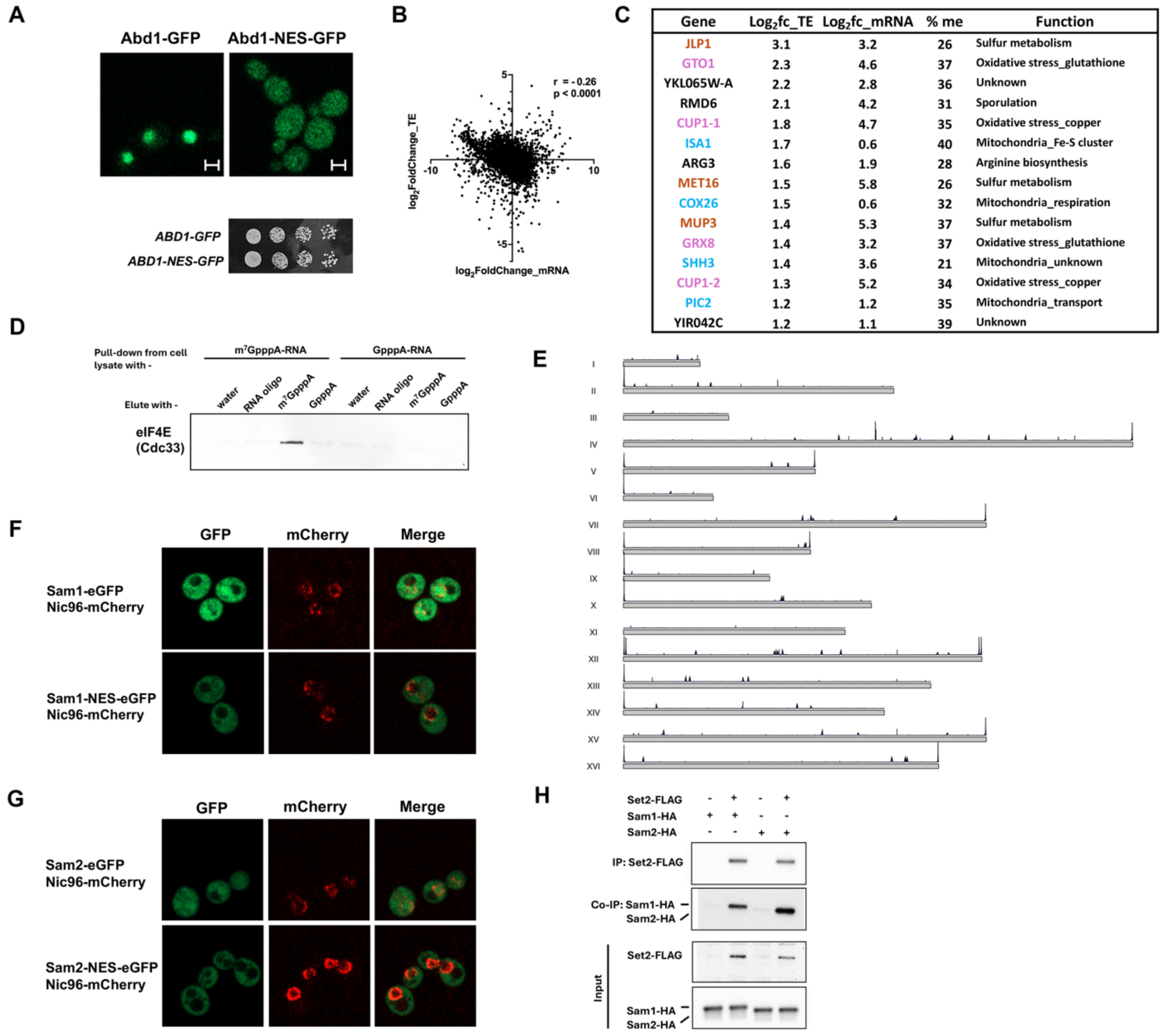


**Fig. S6. (A)** Microscopy images of yeast strains expressing Abd1-GFP and Abd1-NES-GFP grown in rich medium, accompanied by growth assay of ABD1-GFP and Abd1-NES-GFP strains on YPL (rich) agar plates. Scale bar shows 2 µm. **(B)** Correlation between $\log_2$FC in TE vs. $\log_2$FC in mRNA levels. Pearson r = -0.26. **(C)** mRNAs with low cap methylation (% methylation (me) < 40), increased expression ($\log_2$FC > 0), and enhanced translation efficiency ($\log_2$FC in TE > 1) after switching from rich to minimal medium. Orange, pink, and blue highlight genes involved in sulfur metabolism, oxidative stress response, and mitochondrial function, respectively. **(D)** Biotinylated RNA pull-down using synthetic RNA oligonucleotides and lysates from the strain with HA-tagged Cdc33 (eIF4E). Bound proteins were eluted with water, an uncapped RNA oligonucleotide, or the $m^7$GpppA or GpppA cap analogs. Cdc33-HA was detected by western blotting. **(E)** Chromosome coverage plot showing distribution of H3K36me3 peaks across chromosomes in cells grown in minimal medium for 1 h. **(F, G)** Fluorescence microscopy images of GFP-tagged Sam1 or Sam2 with or without a nuclear export signal (NES). The nuclear envelope was labeled with Nic96-mCherry. **(H)** Co-immunoprecipitation of Set2-FLAG with Sam1-HA or Sam2-HA. Set2-FLAG was immunoprecipitated using anti-FLAG M2 antibody. Set2-untagged strains were used as negative controls.
